# Inactivation of the RB1 and PTPN14 tumor suppressors cooperatively enables the lifespan extension activity of the human papillomavirus E7 oncoprotein

**DOI:** 10.64898/2026.03.16.712171

**Authors:** Pushkal Sinduvadi Ramesh, Angelo A. Nicolaci, Leah E. Graham, Joangela Nouel, Kevin Xu, Jennifer M. Binning, Karl Munger, Elizabeth A. White

## Abstract

High-risk human papillomavirus (HPV) E7 proteins bind and inactivate host cellular tumor suppressors and are essential for the immortalization of primary human keratinocytes. E7 proteins from high- and low-risk HPV genotypes bind directly to at least two tumor suppressors, RB1 and PTPN14, and inactivate both. We previously characterized mutations in high-risk HPV E7 proteins that selectively abrogate the ability of E7 to bind either RB1 or PTPN14. Here, we established a genetic complementation system using the E7 mutants defective for binding to RB1 or PTPN14. Neither mutant alone could extend the lifespan of primary keratinocytes. When expressed together, the mutants could, like wild-type high-risk HPV E7, extend keratinocyte lifespan. Both high- and low-risk E7 reduced PTPN14 protein levels and reduced expression of keratinocyte differentiation genes, whereas only high-risk E7 reduced steady-state RB1 levels and induced E2F-dependent genes. Depletion of either RB1 or PTPN14 could cooperate with low-risk HPV6 E7 to extend keratinocyte lifespan. Our findings advance the model that inactivation of at least two tumor suppressors is required for the lifespan extension activity of high-risk HPV E7.

**Significance:** Inactivation of the retinoblastoma tumor suppressor (RB1) is necessary but insufficient for HPV E7-mediated immortalization of human cells. In addition to inactivating RB1, HPV E7 proteins also target for degradation PTPN14, a tumor suppressor and inhibitor of the YAP1 oncoprotein. We report genetic complementation experiments demonstrating that RB1 inactivation and PTPN14 inactivation are separate activities of E7. Either depletion of RB1 or PTPN14 can confer lifespan extension activity on a low-risk HPV E7. These findings redefine our understanding of E7 transforming activity. The predominant difference between high- and low-risk E7 is their ability to degrade RB1, but inactivation of both tumor suppressors is required for E7 activity. Targeting either E7/RB1 or E7/PTPN14 could be of therapeutic benefit.

## Introduction

Human papillomaviruses (HPVs) are double-stranded DNA viruses that infect cutaneous or mucosal epithelial tissues, and several hundred genetically distinct HPV genotypes have been identified (1). Most HPV infections are transient and asymptomatic. However, persistent infection with a subset of high-risk (oncogenic) HPVs can cause mucosal epithelial cancers including cervical, other anogenital, and oropharyngeal cancers (2–4). Low-risk HPVs also infect mucosal tissues but do not cause cancer in healthy individuals at the same anatomical sites.

HPV16 and HPV18 are the most prevalent high-risk HPV genotypes worldwide. They cause most cervical cancers and HPV-associated oropharyngeal cancers in many regions of the world (5). HPV-mediated carcinogenesis is driven by the E6 and E7 oncoproteins, which inactivate cellular tumor suppressor proteins (6). Cellular immortalization is a necessary precursor to carcinogenesis, and the combined expression of high-risk HPV E6 and E7 is necessary and sufficient to immortalize otherwise short-lived primary epithelial cells in culture (7–9). High-risk HPV E7 alone can extend the lifespan of these cells, i.e., enable their growth for several months. Keratinocyte lifespan extension is not a characteristic of low-risk E7 or of high-risk or low-risk E6 proteins (10, 11).

HPV E7 proteins alter cellular signaling by binding to host proteins. Although high- and low-risk HPV E7 proteins share several host cell targets, they have different carcinogenic potential (12, 13). Two of the host cell proteins that are bound by HPV E7 from many different genotypes are the tumor suppressors retinoblastoma protein (RB1) and protein tyrosine phosphatase non-receptor type 14 (PTPN14) (12, 14–17). The crystal structures of the E7-RB1 and E7-PTPN14 interaction sites have been solved, demonstrating that both RB1 and PTPN14 bind directly to E7 (18, 19). E7 binding to either RB1 or PTPN14 requires amino acids in E7 that are highly conserved across the HPV phylogeny (14, 15, 20, 21). Although many other binding partners of E7 have been reported, we are unaware of other E7-host protein interactions that occur via direct binding to similarly highly conserved sites.

HPV E7 proteins bind to RB1 using an LxCxE amino acid sequence in E7 conserved region 2 (CR2) (14, 15). HPV E7 binds predominantly to hypophosphorylated RB1, which is the species that can bind and inhibit E2F transcription factors (22–24). High- and low-risk E7 have different effects on RB1. High-risk HPV E7 proteins are more efficient than low-risk HPV E7 proteins in both their binding to RB1 and their disruption of the E2F/RB1 complex (14, 25–30). High-risk HPV E7 proteins also target RB1 for degradation, as measured by a decrease in RB1 half-life (31, 32). One consequence of disrupting RB1/E2F complexes is increased transcription of genes required for cell cycle progression from G1 into S phase (33). However, there are differing reports regarding the magnitude of E2F target gene induction by E7. In reporter assays in human, primate, or rodent cell lines, high-risk HPV E7 proteins cause a robust increase in E2F reporter transcription (34–37). In contrast, the effect of low-risk HPV E7 on E2F reporters varies in the same experiments. We are unaware of experiments that have compared the effect of high- vs low-risk HPV E7 on endogenous E2F target genes in human keratinocytes, and consequently the question of whether high- and low-risk HPV E7 have equivalent capacity to induce E2F dependent gene expression remains unresolved.

Like RB1, PTPN14 binds to E7 proteins from many different HPV genotypes (12, 21). Work from several groups has established the mechanism by which HPV E7 proteins bind PTPN14 and target it for degradation. The N-terminus of E7 binds to the ubiquitin ligase UBR4/p600 and the C-terminus binds to PTPN14, forming an E7/PTPN14/UBR4 complex and enabling PTPN14 ubiquitination and degradation (38–40). The structure of the HPV18 E7 C-terminus bound to the C-terminus of PTPN14 has been solved, revealing that HPV18 E7 R84 and L91 make direct contact with PTPN14 via intermolecular hydrogen bonds and hydrophobic interactions, respectively (18). The amino acids in PTPN14 that bind to E7 are required for PTPN14 to inhibit the growth of HeLa cells in soft agar (18). The C-terminal arginine that corresponds to R84 in HPV18 E7 (and R77 in HPV16 E7) is highly conserved (21). Of the 219 HPV genotypes in genus alpha, beta, and gamma, there is an arginine at the same site in 84.5% of the E7 proteins and another basic amino acid (lysine or histidine) at the same site in nearly every other E7. The amino acids in the region of E7 that binds to UBR4 are likewise highly conserved (21).

PTPN14 is a tumor suppressor that negatively regulates the YAP1 oncoprotein, which is a driver of cell stemness and self-renewal (41–47). PTPN14 mutations occur frequently in several cancer types (48, 49). We found that when HPV16 or HPV18 E7 degrades PTPN14, YAP1 becomes active and localizes to the nucleus (50–54). In monolayer cell culture, the primary effect of PTPN14 degradation is decreased expression of early keratinocyte differentiation markers such as keratins 1 and 10 (55). PTPN14 requires YAP1 to regulate differentiation genes (50). Consistent with a reduced commitment to epithelial differentiation, when E7 can degrade PTPN14, keratinocytes are preferentially retained in the basal layer of a stratified epithelium (50). PTPN14 likely lacks phosphatase activity. It does not contain a sequence that, in active protein tyrosine phosphatases, contributes to binding phosphotyrosine, nor does mutation of the putative PTPN14 active site prevent PTPN14 from relocalizing YAP1 to the cytoplasm or restricting cell growth (44, 49, 56). The tumor suppressive capacity of PTPN14 instead depends on its PPxY motifs, which bind to cellular proteins to promote activity of the Hippo pathway upstream of YAP1 (49, 51).

Here, we sought to determine whether and how RB1 inactivation and PTPN14 inactivation cooperate to enable certain biological activities of high- and/or low-risk HPV E7. We focused here on how E7 reprograms host cell transcription and how the resulting transcriptional changes correlate with the ability of E7 to extend the lifespan of primary keratinocytes. Lifespan extension is an essential precursor to carcinogenesis. Our experiments address several questions. One question is related to RB1-independent activities of E7. RB1 binding and/or degradation is necessary, but insufficient, for high-risk HPV E7 carcinogenic activity (27, 57–65). Consequently, one or more RB1-independent activities must cooperate with RB1 inactivation to enable transformation by E7, but there is no consensus as to the nature of such an activity. PTPN14 inactivation could be an RB1-independent transforming activity. We and others established that the E7/PTPN14/UBR4 complex can be separated by size-exclusion chromatography from the E7/RB1 complex, that mutation of the RB1, PTPN14, and/or UBR4 binding sites reduces E7 carcinogenic activity, and that many of the genes dysregulated upon PTPN14 inactivation are different from those dysregulated by HPV E7 binding to RB1 (17, 40, 55). However, we lacked genetic evidence in support of two independent activities of HPV E7. A second question is why high- and low-risk HPV E7 proteins share many cellular targets yet only high-risk E7 can transform. Here we addressed both questions. We demonstrate genetic complementation using RB1- and PTPN14-binding deficient mutants of HPV16 or HPV18 E7 in keratinocyte lifespan extension assays. Comparing the effect of high- and low- risk HPV E7 on PTPN14 and RB1, we found that various E7s shared the ability to reduce PTPN14 levels and inhibit differentiation gene expression, but only high-risk E7 caused a robust increase in E2F-dependent gene expression downstream of RB1 inactivation. Either RB1 knockdown or PTPN14 knockdown cooperated with low-risk HPV E7 in assays of keratinocyte lifespan extension. We conclude that inactivation of both RB1 and PTPN14 is required for the lifespan extension activity of high-risk HPV E7, and experimentally induced loss of either RB1 or PTPN14 enables lifespan extension in the presence of low-risk E7.

## Results

### High-risk HPV E7 mutants defective for RB1 or PTPN14 binding complement in *trans* to extend keratinocyte lifespan

We hypothesized that if RB1 inactivation and PTPN14 inactivation are independent activities of high-risk HPV E7, then in assays of keratinocyte lifespan extension a mutant that cannot inactivate RB1 would complement one that cannot inactivate PTPN14. To enable genetic complementation experiments, we used pairs of retroviral vectors with complementary selectable markers to deliver two different versions of E7 to the same cell. First, we focused on HPV18 E7. To disrupt binding of E7 to RB1, we used HPV18 E7 ΔDLLC, which harbors a deletion in the LxCxE motif required for E7 to bind RB1 (24, 50). To disrupt binding of E7 to PTPN14, we used HPV18 E7 R84A L91A, which was previously characterized by Yun and colleagues and is altered in two residues that make direct contact with PTPN14 (18). Like HPV18 E7 R84S, which we previously characterized (21), HPV18 E7 R84A L91A was impaired in reducing the steady-state levels of PTPN14 protein but competent to induce E2F target genes PCNA and CCNE2 (Supplemental Figure 1A-B).

To confirm that RB1 and PTPN14 binding defective HPV18 E7 mutants were impaired in keratinocyte lifespan extension assays, we expressed wild-type HPV18 E7, HPV18 E7 ΔDLLC, or HPV18 E7 R84A L91A, or GFP as a negative control, in primary human foreskin keratinocytes (HFKs) (Supplemental Figure 1C-E). HPV18 E7 ΔDLLC (cannot bind RB1) was delivered using a vector conferring resistance to hygromycin whereas HPV18 E7 R84A L91A (cannot bind PTPN14) was delivered using a vector conferring resistance to neomycin. Cells expressing wild-type HPV18 E7 exhibited reduced steady-state levels of RB1 and PTPN14 proteins, increased expression of E2F target genes PCNA and CCNE2, and decreased expression of early differentiation markers KRT1 and KRT10 compared to the GFP control. HPV18 E7 ΔDLLC reduced the steady-state level of PTPN14, but not RB1, and did not induce the expression of PCNA or CCNE2. HPV18 E7 R84A L91A reduced the steady state level of RB1 and induced PCNA and CCNE2 but did not reduce PTPN14 levels. KRT1 and KRT10 expression correlated with the level of PTPN14 in the cells. Neither mutant could promote cell growth comparable to wild-type HPV18 E7 (Supplemental Figure 1F). HPV18 E7 ΔDLLC was unable to extend keratinocyte lifespan (as assessed by time in culture) or promote proliferation (as assessed by cumulative population doublings). A few HPV18 E7 R84A L91A cells were maintained for 50 days, but their proliferative capacity was low throughout the experiment.

Next, we tested whether the two mutants could complement one another in *trans*. Primary HFKs were transduced with pairs of retroviruses encoding GFP, wild-type HPV18 E7, HPV18 E7 ΔDLLC, or HPV18 E7 R84A L91A, and selected with hygromycin and neomycin. Combined expression of HPV18 E7 ΔDLLC and HPV18 E7 R84A L91A reduced the steady-state levels of RB1 and PTPN14 (Figure 1A), induced the expression of E2F targets PCNA and CCNE2 (Figure 1B), and inhibited the expression of early differentiation markers KRT1 and KRT10 (Figure 1C). In each case, the combined expression of the two mutant proteins recapitulated the effect of wild-type HPV18 E7. In the keratinocyte lifespan extension assay, cells transduced with HPV18 E7 ΔDLLC (cannot bind RB1) doubled only as many times as the GFP control cells (Figure 1D). Cells transduced with HPV18 E7 R84A L91A (cannot bind PTPN14) could be maintained in culture for up to 120 days, but the cells formed colonies poorly and doubled only ∼20 times over the three-month assay. In contrast, the combined expression of HPV18 E7 ΔDLLC plus HPV18 E7 R84A L91A enabled HFK growth. The cells transduced with both mutants were, like those expressing wild-type HPV18 E7, readily maintained in culture for up to 120 days. The cells expressing wild-type HPV18 E7 doubled 52 times and the cells expressing both mutants doubled 44 times during the assay.

**Figure 1:**
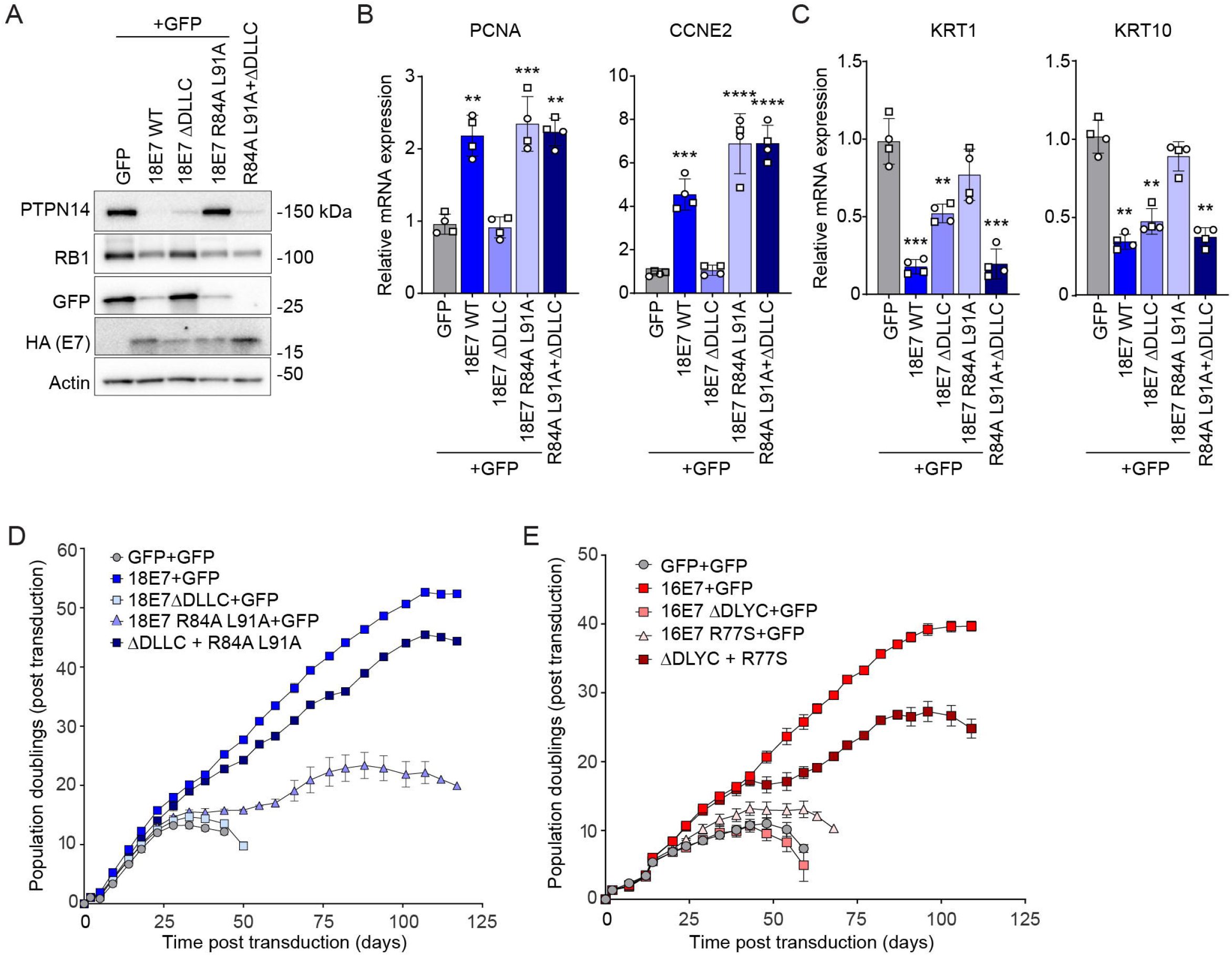
High-risk HPV E7 mutants defective for RB1 or PTPN14 binding complement in *trans* to extend keratinocyte lifespan. (A-D) Primary human keratinocytes were transduced with pairs of retroviruses encoding GFP, wild-type HPV18 E7 (WT), HPV18 E7 ΔDLLC, or HPV18 E7 R84A L91A, with one hygromycin-resistant and one neomycin-resistant retrovirus included in each condition. Each version of HPV18 E7 has an HA epitope tag at its C-terminus. (A) Total cell lysates were subjected to western blotting and probed with antibodies to PTPN14, RB1, GFP, HA, and actin. The expression of (B) E2F target genes (PCNA, CCNE2) and (C) differentiation markers (KRT1, KRT10) were measured by qRT-PCR. Bar graphs represent the mean ± SD of two independent experiments each conducted in duplicate. Statistical significance was determined by one-way ANOVA followed by Holm-Sidak multiple comparison test, comparing each condition to the GFP control (**=*p* ≤ 0.01; ***=*p* ≤ 0.001; ****=*p* ≤ 0.0001). (D) Cell populations were cultured for up to 117 days. Cells were counted at each passage and cell count data was used to calculate cumulative population doublings. Data points indicate the mean and error bars indicate the range for two replicate cell populations. Error bars that are not shown are not visible at this scale. (E) Primary human keratinocytes were transduced with pairs of retroviruses encoding GFP, wild-type HPV16 E7 (WT), HPV16 E7 ΔDLYC, or HPV16 E7 R77S. One hygromycin-resistant and one neomycin-resistant retrovirus was included in each condition. Each version of HPV16 E7 has an HA epitope tag at its C-terminus. Cell populations were cultured for up to 109 days. Cells were counted at each passage and cell count data was used to calculate cumulative population doublings. Data points indicate the mean and error bars indicate the range for two replicate cell populations.

We repeated the complementation analysis for HPV16 E7, first confirming that delivering the HPV16 E7 mutants with hygromycin or neomycin resistant vectors had the same effects as the puromycin resistant vectors we previously used (21, 55). Like HPV18 E7, wild-type HPV16 E7 induced the expression of E2F targets PCNA and CCNE2 and reduced the expression of early differentiation markers KRT1 and KRT10 (Supplemental Figure 2A, C). HPV16 E7 ΔDLYC (cannot bind RB1) did not induce E2F target expression but impaired the expression of early differentiation markers. HPV16 E7 R77S (cannot bind PTPN14) induced E2F-dependent gene expression but did not inhibit the expression of early differentiation markers (Supplemental Figure 2A, C). In lifespan extension assays lasting 60-70 days, cells transduced with wild-type HPV16 E7 retrovirus doubled 20-25 times, whereas cells transduced with a single retrovirus expressing HPV16 E7 ΔDLYC or HPV16 E7 R77S doubled fewer times and/or could not be maintained in culture for more than 60 days (Supplemental Figure 2B, D). Combined expression of the two defective mutants recapitulated the effect of wild-type HPV16 E7 on PTPN14 and RB1 protein levels and expression of E2F target and early differentiation marker genes (Supplemental Figure 3).

We then tested the ability of the individual HPV16 E7 mutants to complement one another in the lifespan extension assay. Similar to HPV18 E7, cells transduced with one HPV16 E7 mutant retrovirus attained at most ∼12 population doublings and stopped proliferating by 75 days (Figure 1E). In contrast, cells transduced with HPV16 E7 ΔDLYC and HPV16 E7 R77S together doubled more than 25 times and, like cells transduced with wild-type HPV16 E7, were readily maintained in culture for approximately four months. Collectively, these findings support that high-risk HPV E7 proteins must bind and degrade both RB1 and PTPN14 in order to extend keratinocyte lifespan. The two activities can be supplied in *trans*, on different molecules of E7.

### High- and low-risk HPV E7 proteins have different effects on RB1 but similar effects on PTPN14

Both the LxCxE motif and the C-terminal arginine are highly conserved among high-and low-risk HPV E7 proteins. However, high-risk HPV E7 proteins can extend the lifespan of primary HFKs whereas low-risk E7 cannot (11). First, we confirmed that high-risk HPV18 E7 but not low-risk HPV6 E7 could promote proliferation and extend primary keratinocyte lifespan in our assay (Figure 2A). We then compared the effects of two high-risk (HPV16, HPV18) and two low-risk (HPV6, HPV11) E7 on the steady-state levels of RB1 and PTPN14. Consistent with earlier literature (66, 67), high-risk HPV E7 reduced the steady-state levels of RB1 and low-risk E7 did not (Figure 2B). There was a corresponding increase in E2F target gene expression in the cells harboring high-risk, but not low-risk HPV E7 (Figure 2C). In contrast, each HPV E7 reduced the steady-state levels of PTPN14 and the expression of early differentiation markers KRT1 and KRT10 (Figure 2B, D).

**Figure 2:**
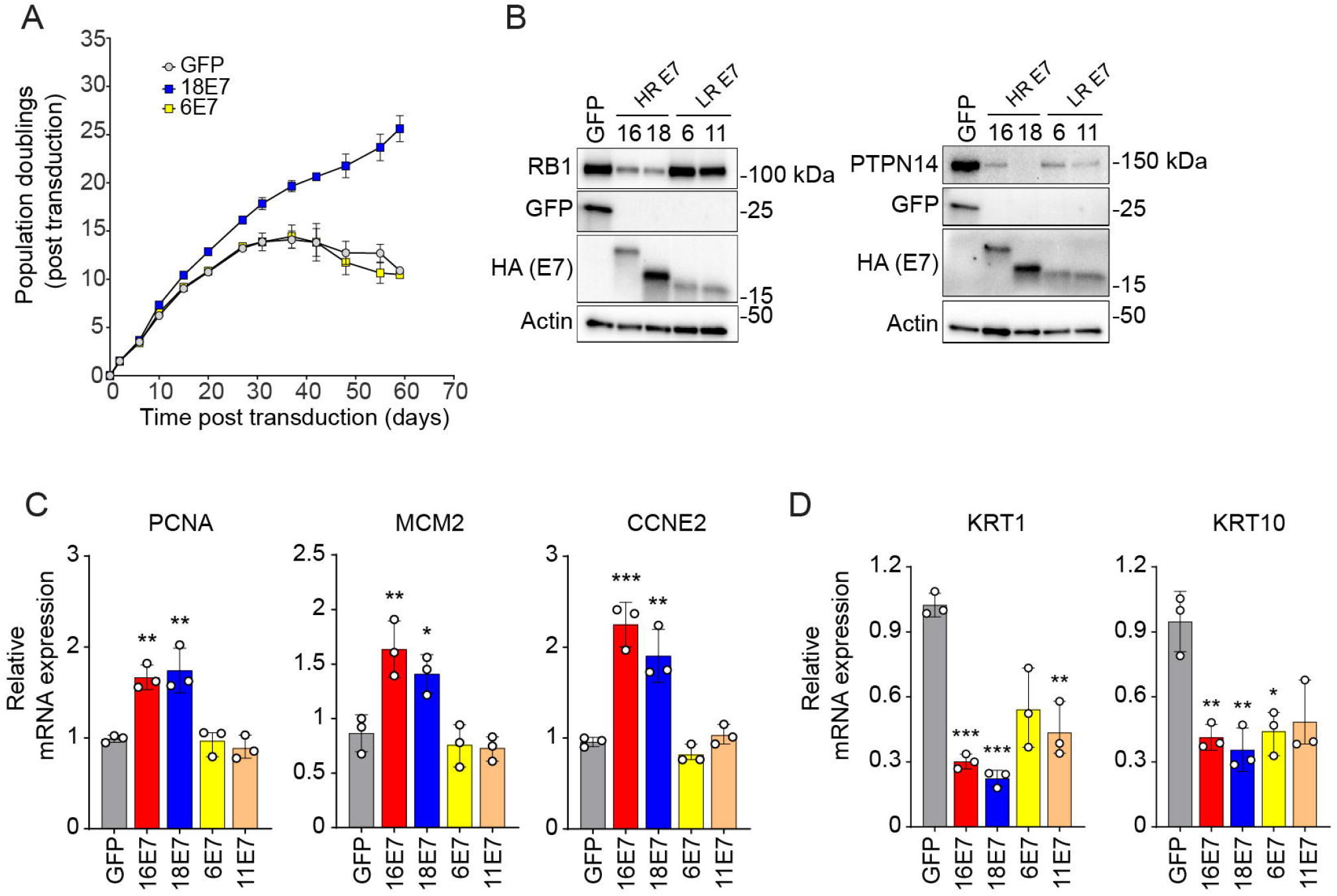
Effect of high-risk and low-risk HPV E7 proteins on RB1 and PTPN14 proteins and target genes. Primary human keratinocytes were transduced with retroviral vectors encoding GFP, HPV16 E7, HPV18 E7, HPV6 E7, or HPV11 E7 and selected with puromycin. Each version of HPV E7 has an HA epitope tag at its C-terminus. (A) Cell pools expressing GFP, HPV18 E7, or HPV6 E7 were cultured for up to 59 days. Cells were counted at each passage and cell count data was used to calculate cumulative population doublings. Data points indicate the mean and error bars indicate the range for two replicate cell populations. (B) Total cell lysates from two experiments were subjected to western blotting and probed with antibodies to PTPN14, RB1, GFP, HA, and actin. HR = high-risk HPV E7; LR = low-risk HPV E7. The expression of (C) E2F targets (PCNA, MCM2, CCNE2) and (D) differentiation markers (KRT1, KRT10) was measured by qRT-PCR. Bar graphs represent the mean ± SD of three replicates. Statistical significance was determined by one-way ANOVA followed by Holm-Sidak multiple comparison test, comparing each condition to the GFP control (*=*p* ≤ 0.05; **=*p* ≤ 0.01; ***=*p* ≤ 0.001).

### HPV16 E7 and HPV6 E7 share the ability to inhibit differentiation gene expression but not the ability to induce E2F target gene expression

Next, we sought to more broadly compare the effect of low- vs high-risk HPV E7 on gene expression. We generated HFKs expressing untagged wild-type high-risk HPV16, low-risk HPV6 E7, or an LxCxE deletion mutant form of each one, validated E7 expression by qRT-PCR (Supplemental Figure 4A), and conducted RNA-seq analysis (Figure 3, Supplemental Figures 4B, 4C, 5). There were 690 differentially expressed genes (≥ 1.5-fold change and adjusted *p-*value ≤ 0.05) in HFK-16E7 compared to HFK-GFP cells (Supplemental Figure 4B). The KEGG pathways enriched among the upregulated genes included those related to the cell cycle, DNA replication, and DNA repair, a signature that is consistent with HPV16 E7-dependent activation of E2F target genes (Figure 3A, Supplemental Table 1). The KEGG pathways enriched among the downregulated genes included several pathways related to keratinocyte differentiation (e.g. cornified envelope formation, integrin signaling, and cadherin signaling), which is consistent with HPV16 E7-dependent degradation of PTPN14 and inhibition of epithelial differentiation. We compared the differentially expressed genes to additional gene sets and found that 38% of the hallmark E2F gene list (GSEA gene set M5925, HALLMARK_E2F_TARGETS), 29% of the PTPN14 knockout signature genes defined in (55), and 60% of the genes differentially expressed in the comparison of HPV16 E7 to HPV16 E7 ΔDLYC were also altered by HPV16 E7 when compared to GFP (Supplemental Figure 4B, 5A, 5B).

**Figure 3:**
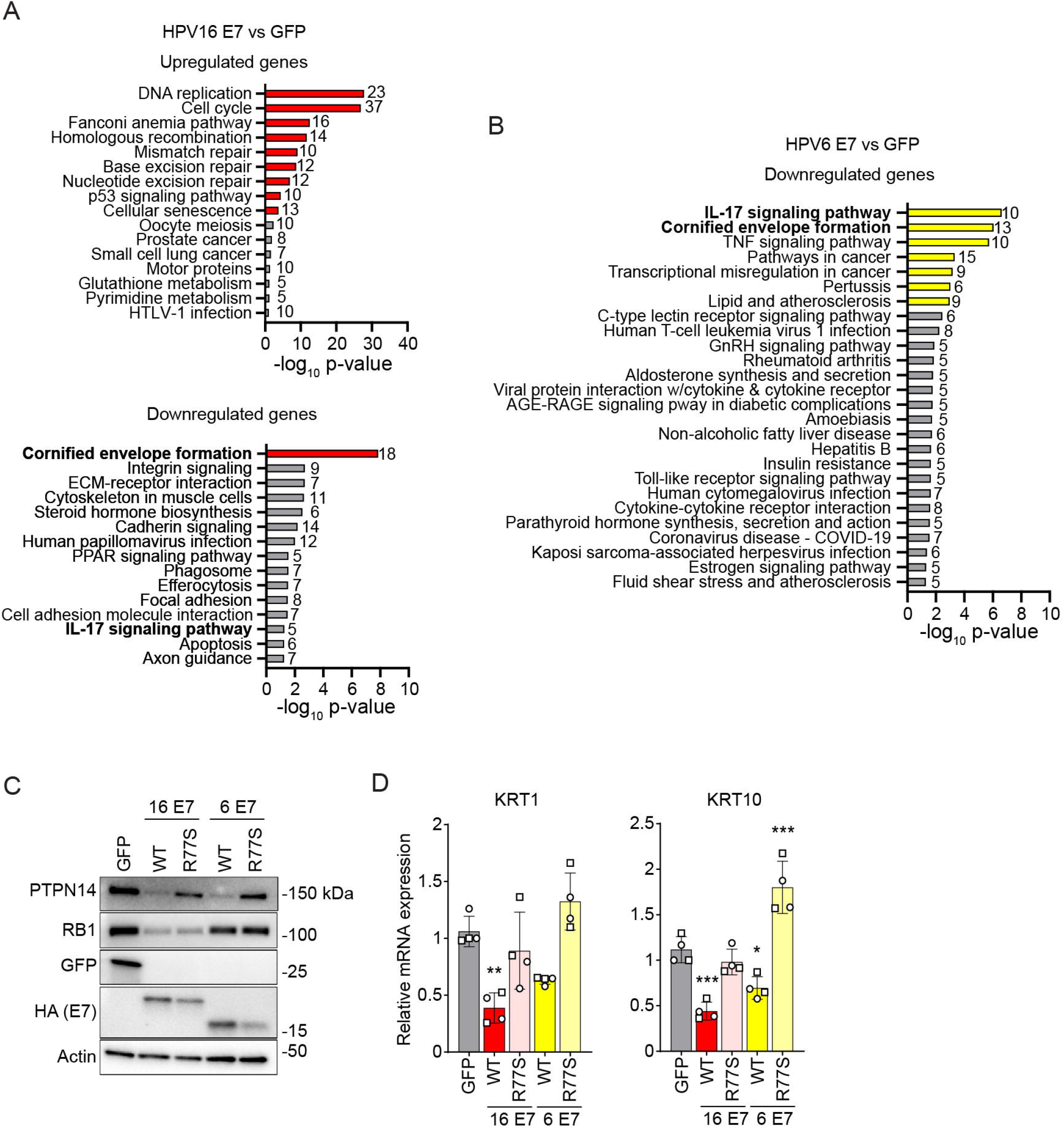
Shared and distinct gene expression patterns in cells expressing high- or low-risk HPV E7. (A, B) Primary HFK were transduced with MSCV retroviral vectors encoding untagged HPV16 or HPV6 E7 proteins or GFP. Following RNA-seq, KEGG pathway enrichment analysis was conducted on the genes that were upregulated (≥1.5-fold higher expression and adjusted *p*-value ≤ 0.05) or downregulated (≥1.5-fold lower expression and adjusted *p*-value ≤ 0.05) in (A) HPV16 E7 cells relative to GFP cells or (B) HPV6 E7 cells relative to GFP cells. Pathways with *p*-values ≤ 0.05 are included in the chart. Bars for pathways with FDR ≤ 0.05 are colored whereas bars for pathways with FDR > 0.05 are in grey. Values to the right of each bar indicate the number of genes in the list associated with a pathway. Pathway names in bold are altered in both comparisons. (C, D) Primary human keratinocytes were transduced with retroviral vectors encoding GFP, wild-type (WT) HPV16 E7, HPV16 E7 R77S, WT HPV6 E7, or HPV6 E7 R77S, and selected with puromycin. Each version of HPV E7 has Flag and HA epitope tags at its C-terminus. (C) Total cell lysates were subjected to western blotting and probed with antibodies to PTPN14, RB1, GFP, HA, and actin. (D) The expression of differentiation markers (KRT1, KRT10) was measured by qRT-PCR. Bar graphs represent the mean ± SD of two independent experiments each conducted in duplicate. Statistical significance was determined by one-way ANOVA followed by the Holm-Sidak multiple comparison test, comparing each condition to the GFP control (*=*p* ≤ 0.05; **=*p* ≤ 0.01; ***=*p* ≤ 0.001).

In contrast, only about one third as many genes (197 genes) were differentially expressed in HFK-6E7 vs HFK-GFP (Supplemental Figure 4C). Unlike HPV16 E7, HPV6 E7 induced the expression of very few genes, and no KEGG pathways were significantly represented among those upregulated genes. Only one gene from the hallmark E2F gene list (CDKN2C) and 7% of the genes differentially expressed in the HPV16 E7 versus the HPV16 E7 ΔDLYC comparison were differentially regulated by HPV6 E7 (Supplemental Figure 4C). In addition, only three genes (SERF1B, APOE, and PI3) were differentially expressed when we compared HPV6 E7 to HPV6 E7 ΔGLHC (cannot bind RB1) (Supplemental Figure 5C) and none of the three were present in the list of hallmark E2F genes. However, similar to HPV16 E7, low-risk HPV6 E7 reduced the expression of genes associated with epithelial differentiation, and the expression of genes in a KEGG pathway related to cornified envelope formation was reduced upon HPV6 E7 expression (Figure 3B). HPV16 E7 and HPV6 E7 altered the expression of a similar number of the PTPN14 knockout signature genes: 29% for HPV16 E7 and 38% for HPV6 E7 (Supplemental Figure 4B, 4C). We conclude that in proliferative cells, HPV6 E7 inhibited differentiation gene expression but did not induce robust E2F target expression.

### HPV6 E7 reduces differentiation gene expression dependent on the C-terminal arginine

Having observed that HPV16 E7 and HPV6 E7 reduced differentiation gene expression to a comparable degree, we tested whether the C-terminal arginine in HPV6 E7 contributes to differentiation repression. We previously determined that wild-type HPV6 E7 reduces the steady-state levels of PTPN14 but HPV6 E7 R77S does not (21). We transduced primary keratinocytes with retroviruses encoding GFP, HPV16 E7, HPV16 E7 R77S, HPV6 E7, or HPV6 E7 R77S. Steady-state levels of PTPN14 protein were reduced in cells expressing wild-type but not R77S mutants of each E7 (Figure 3C). Consistent with our previous finding that HPV16 E7 and HPV18 E7 had a greater ability to reduce KRT1 and KRT10 protein levels in differentiated cells than their arginine mutant counterparts (21), *KRT1* and *KRT10* mRNA levels were lower in cells expressing the wild-type form of each E7 compared to the corresponding R77S mutant (Figure 3D). These findings indicate that inhibition of differentiation gene expression by either high- or low-risk E7 depends on the conserved C-terminal arginine.

### HPV18 E7 binds to PTPN14 with a higher affinity than HPV16, 6, or 11 E7

Having observed that high- and low-risk HPV E7 share the ability to reduce PTPN14 levels and differentiation gene expression, we sought to determine the affinity of the same E7 proteins for the C-terminal phosphatase domain of PTPN14. Full-length HPV16, 18, 6, or 11 E7 were purified and used in biolayer interferometry (BLI) experiments together with the purified C-terminal phosphatase domain of PTPN14 (Figure 4A, Supplemental Figure 6). The dissociation constants were determined for each pairwise interaction between an E7 and PTPN14 (Figure 4B). The affinity of E7 for PTPN14 was highest for HPV18 E7 (*K*_D_ = 26.6 ± 12.8 nM), next highest for HPV16 E7 (*K*_D_ = 112.5 ± 33.7), and lower for the two low-risk HPV6 and HPV11 E7 (*K*_D_ = 189.5 ± 58.7, and 187.9 ± 22.7 nM, respectively).

**Figure 4:**
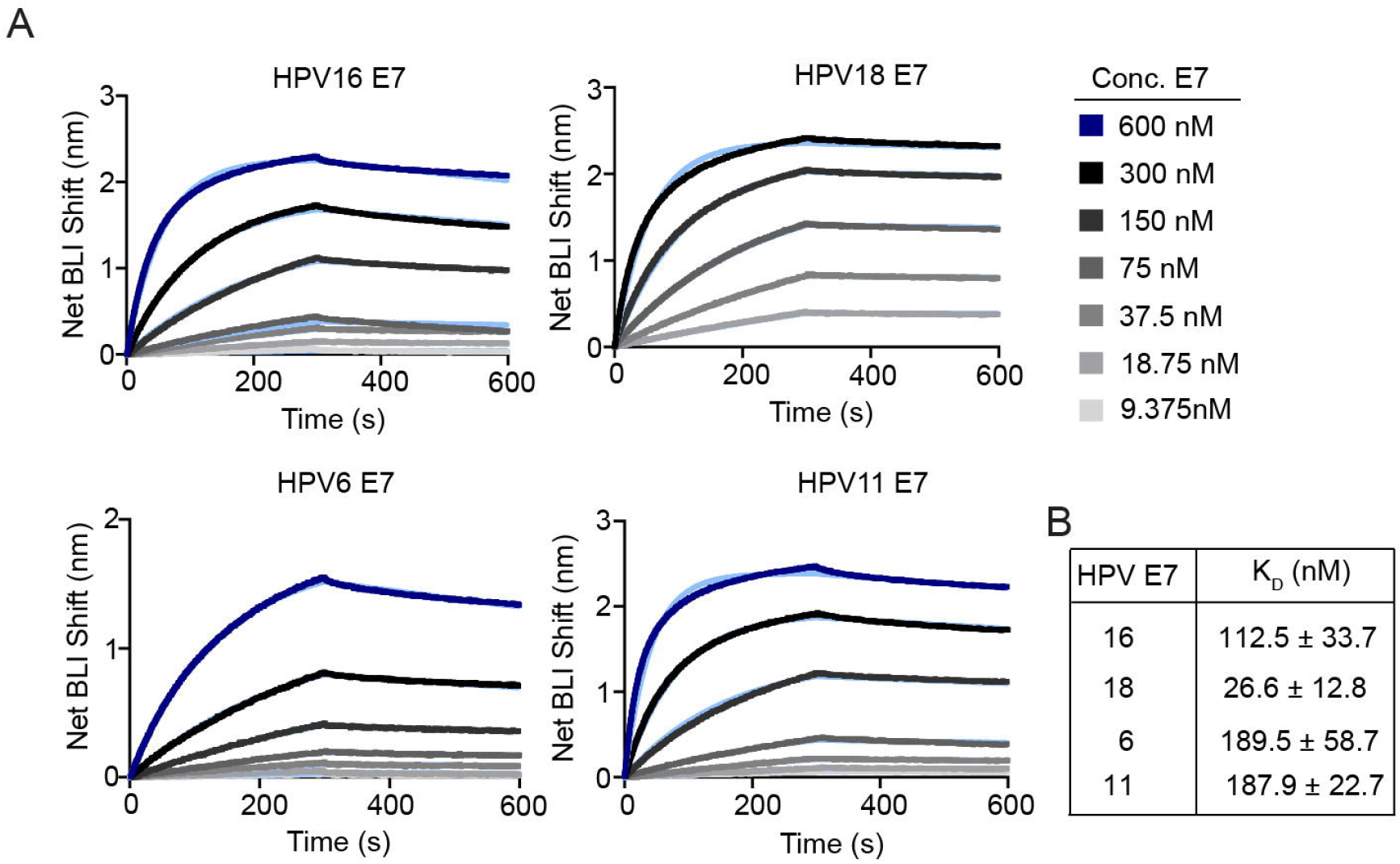
BLI analysis of PTPN14 binding to high- or low-risk HPV E7 proteins. (A) Affinity measurements of twin-strep tagged phosphatase domain of PTPN14 (amino acids 886-1147) against MBP-HPV E7 proteins from four genotypes were conducted using bio-layer interferometry (BLI). Dark blue and black traces represent the raw data; the kinetic fit is shown in light blue underneath. Sensograms shown are representative plots of three replicates for each E7. (B) Table displays equilibrium constants (K_D_) for each interaction.

### HPV6 E7 and RB1 knockdown cooperate to extend keratinocyte lifespan

Having observed that the magnitude of RB1 inactivation distinguishes high-risk from low-risk E7, we hypothesized that knockdown of RB1 would complement a low-risk HPV E7 in a keratinocyte lifespan extension assay. We generated all-in-one CRISPR interference (CRISPRi) lentiviruses to reduce RB1 transcript levels in primary HFKs. The CRISPRi vector encodes dCas9-KRAB-MeCP2 (68) plus one of three non-targeting (sgNT) sgRNAs or one of four sgRNAs directed to the RB1 promoter. Three of the RB1 CRISPRi sgRNAs reduced RB1 protein and RNA levels and induced expression of the E2F target CCNE2 (Figure 5A, B). We transduced primary HFKs with a pair of vectors: the CRISPRi sgNT or sgRB1 vector plus a second vector encoding GFP, high-risk HPV18 E7, or low-risk HPV6 E7. RB1 depletion reduced RB1 protein and RNA levels and induced CCNE2 in the presence or absence of HPV6 E7 (Figure 5C, D). Neither HPV6 E7 nor RB1 depletion alone extended the lifespan of HFKs compared to the GFP control (Figure 5E). In contrast, combined expression of HPV6 E7 and depletion of RB1 enabled long-term propagation of the cells. Like the HPV18 E7 cells, the HPV6 E7 + CRISPRi sgRB1 cells were readily maintained in culture for 90 days, and they doubled up to 20 times during the experiment. To confirm that RB1 depletion alone was insufficient to enable lifespan extension, even in the presence of HPV6 E7, we repeated the experiment, this time combining the PTPN14 binding-deficient HPV6 E7 R77S mutant with RB1 knockdown (Figure 5F, G). Neither RB1 depletion alone nor the combination of sgRB1 and HPV6 E7 R77S expression could extend keratinocyte lifespan. Only the conditions in which HPV6 E7 could bind PTPN14 and RB1 was inactivated enabled keratinocyte culture for longer than 60 days (Figure 5G).

**Figure 5:**
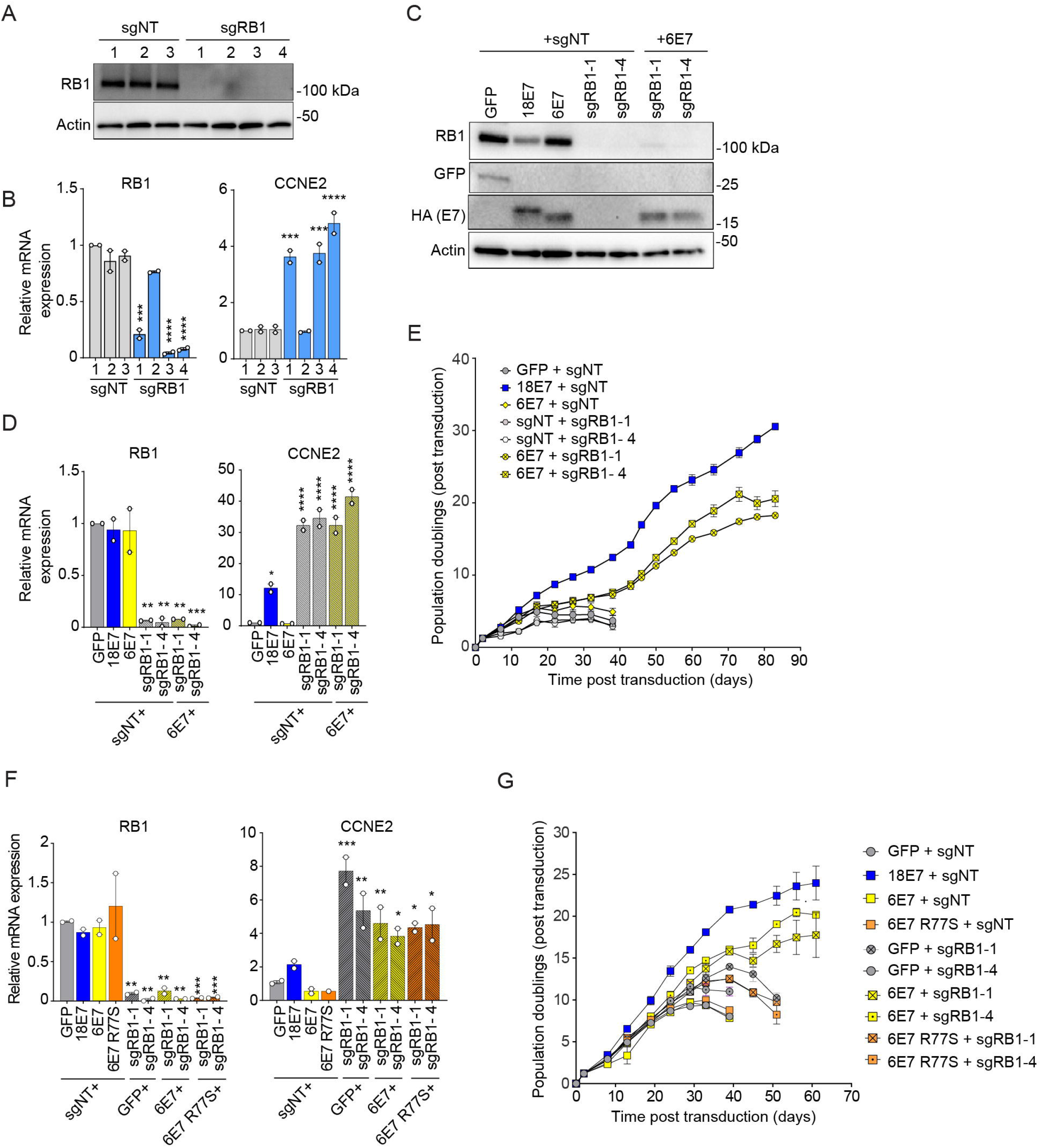
RB1 knockdown promotes lifespan extension in primary keratinocytes expressing HPV6 E7. (A, B) Primary human keratinocytes were transduced with a CRISPRi lentiviral vector encoding dCas9-KRAB MeCP2 and a non-targeting sgRNA or a sgRNA targeting RB1. (A) Total cell lysates were subjected to western blotting and probed with antibodies to RB1 and actin. (B) The expression of RB1 and CCNE2 was measured by qRT-PCR. Bar graphs represent the mean ± range of two replicate cell populations. (C-E) Primary human keratinocytes were transduced with a pair of vectors: one retrovirus encoding GFP, HPV18 E7, or HPV6 E7, plus one CRISPRi lentivirus, and selected with puromycin and neomycin. (C) Total cell lysates were subjected to western blotting and probed with antibodies to RB1, GFP, HA, and actin. (D) The expression of RB1 and CCNE2 was measured by qRT-PCR. (E) Cell populations were cultured for up to 83 days. Cells were counted at each passage and cell count data was used to calculate cumulative population doublings. Data points indicate the mean and error bars indicate the range for two replicate cell populations. (F, G) Primary human keratinocytes were transduced with a pair of vectors: one retrovirus encoding GFP, HPV18 E7, HPV6 E7, or HPV6 E7 R77S plus one CRISPRi lentivirus encoding dCas9-KRAB MeCP2 and a non-targeting sgRNA (sgNT) or a sgRNA targeting RB1 (sgRB1) and selected with puromycin and neomycin. (F) The expression of RB1 and CCNE2 was measured by qRT-PCR. Bar graphs represent the mean ± range of two replicate experiments. Statistical significance was determined by one-way ANOVA followed by the Holm-Sidak multiple comparison test, comparing each condition to the matched sgNT control (*=*p* ≤ 0.05; **=*p* ≤ 0.01; ***=*p* ≤ 0.001; ****=*p* ≤ 0.0001). (G) Cell populations were cultured for up to 60 days. Cells were counted at each passage and cell count data was used to calculate cumulative population doublings. Data points indicate the mean and error bars indicate the range for two replicate cell populations.

### HPV6 E7 and PTPN14 knockdown cooperate to extend keratinocyte lifespan

Considering that the binding affinity assays revealed that high-risk bind to PTPN14 with higher affinity than low risk HPV E7, we next tested whether depletion of PTPN14 would complement a low-risk HPV E7 in a keratinocyte lifespan extension assay. We generated vectors for CRISPRi depletion of PTPN14 and confirmed that they reduced PTPN14 protein and RNA levels and expression of the differentiation marker KRT10 (Figure 6A, B). We transduced primary HFKs with one vector that expressed either GFP control, high-risk HPV16 E7, or low-risk HPV6 E7, plus a second CRISPRi sgNT or sgPTPN14 vector (Figure 6C). PTPN14 knockdown reduced PTPN14 transcript levels but did not further reduce KRT10 expression in HPV16 E7 or HPV6 E7 expressing keratinocytes (Figure 6D). Neither HPV6 E7 nor PTPN14 depletion alone extended keratinocyte lifespan compared to control cells (Figure 6E). In contrast, PTPN14 knockdown cooperated with HPV6 E7 to extend keratinocyte lifespan, and the cells transduced with both vectors were maintained in culture for 116 days. Depleting PTPN14 also increased the growth of HPV16 E7 cells, enabling HPV16 E7 + sgPTPN14 cells to double about five more times than HPV16 E7 + sgNT cells during the four-month assay.

**Figure 6:**
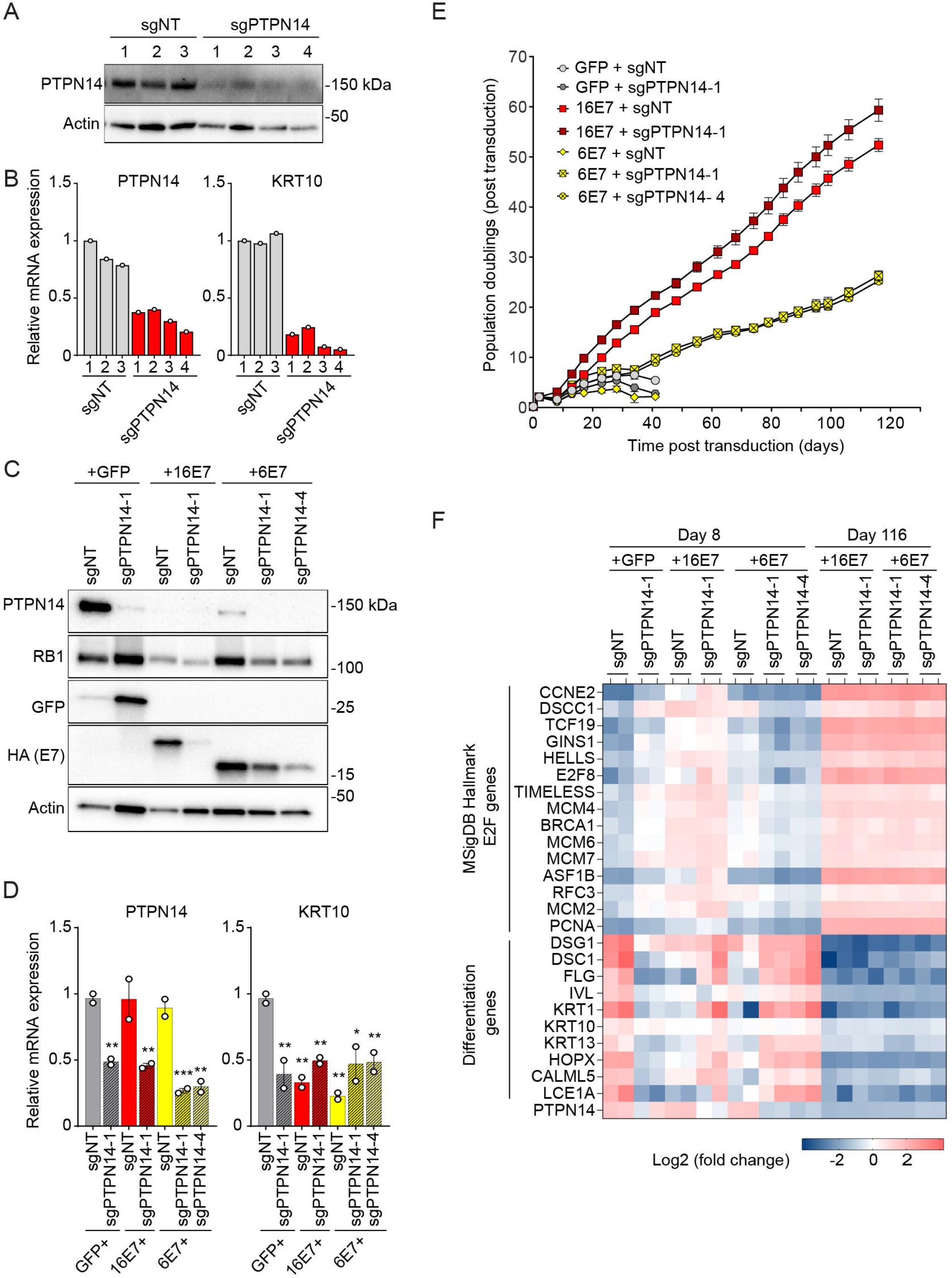
PTPN14 knockdown promotes lifespan extension in primary keratinocytes expressing HPV6 E7. (A, B) Primary human keratinocytes were transduced with a lentiviral CRISPRi vector encoding dCas9-KRAB MeCP2 and a non-targeting sgRNA or a sgRNA targeting PTPN14. (A) Total cell lysates were subjected to western blotting and probed with antibodies to PTPN14 and actin. (B) The expression of PTPN14 and KRT10 was measured by qRT-PCR. (C-E) Primary human keratinocytes were transduced with a combination of one retrovirus encoding GFP, HPV16 E7, or HPV6 E7, plus one CRISPRi lentivirus and selected with puromycin and neomycin. (C) Cell lysates were subjected to western blotting and probed with antibodies to PTPN14, RB1, GFP, HA, and actin. (D) The expression of PTPN14 and KRT10 was measured by qRT-PCR. Bar graphs represent the mean ± range of two replicate cell populations. Statistical significance was determined by one-way ANOVA followed by the Holm-Sidak multiple comparison test, comparing each condition to the GFP control (*=*p* ≤ 0.05; **=*p* ≤ 0.01, ***=*p* ≤ 0.001). (E) Cell populations were cultured for up to 116 days. Cells were counted at each passage and cell count data was used to calculate cumulative population doublings. Data points indicate the mean and error bars indicate the range for two replicate cell populations. (F) Heat map showing the expression of selected MSigDB Hallmark E2F target genes and keratinocyte differentiation genes, as determined by RNA sequencing, in cells collected at day 8 or day 116 of the growth curve in (E). Gene expression values are displayed as row-scaled normalized expression.

To assess whether global patterns of gene expression could explain the observed increase in lifespan and proliferation of the HPV6 E7 + sgPTPN14 cells, we sequenced RNA from all the cell populations early after transduction (Figure 6F, Day 8) and all of the cell populations that remained after several months in culture (Figure 6F, Day 116). At early times post transduction, HPV16 E7 induced the expression of many E2F target genes in the MSig Database list of Hallmark E2F genes, and there were differences in gene expression in HPV16 E7 vs HPV6 E7 cells consistent with those identified in the previous RNA-seq experiment. At late times, HPV16 E7 and HPV6 E7 continued to express the version of E7 with which they were originally transduced (Supplemental Figure 7), but all the cell pools exhibited upregulated expression of E2F target genes and downregulated expression of differentiation related genes compared to any cell pool at the early time point. With respect to E2F target genes and keratinocyte differentiation genes, all of the surviving cell pools converged upon a shared gene expression signature that, for HPV6 E7, was enabled by simultaneous depletion of PTPN14.

## Discussion

Our previous results prompted the hypothesis that RB1 inactivation and PTPN14 inactivation are separate activities of high-risk HPV E7 proteins that are both required for E7 carcinogenic activity. Here, we tested this hypothesis using mutants of high-risk HPV E7 in genetic complementation experiments. When we co-expressed mutants of HPV16 or HPV18 E7, providing the LxCxE motif in one E7 molecule and the PTPN14 binding site in another, the combination was sufficient to recapitulate the effect of wild-type high-risk HPV16 or HPV18 E7 in several assays. In particular, two individually impaired mutants cooperated to extend the lifespan of primary keratinocytes (Figure 1). We note that compared to wild-type E7, combined expression of the two mutants enabled cell propagation for the same length of time but supported somewhat fewer population doublings (Figure 1). Regardless, cells expressing either wild-type E7 or the two mutants together exhibited both lifespan extension (reflected by time in culture) and induction of proliferation (reflected by population doublings), the combination of which was not exhibited by cells expressing either mutant alone. The ability of HPV E7 to extend keratinocyte lifespan over 3-4 months is well correlated with its ability to immortalize keratinocytes in combination with HPV E6 (11), and keratinocyte immortalization is an essential early step in HPV carcinogenesis. We previously showed that a mutant form of HPV16 E7 that cannot bind UBR4 or degrade PTPN14 is impaired in a lifespan extension assay even in the presence of HPV16 E6 (55). Our results support that RB1 inactivation and PTPN14 inactivation are independent activities of high-risk HPV E7 that are both required for keratinocyte immortalization.

High- and low-risk HPV E7 proteins engage several of the same cellular targets but have different biological activities. We sought to determine whether and how different effects on RB1 and/or PTPN14 account for the differences between high- and low-risk HPV E7 proteins. Consistent with previous publications (66, 67), high-risk but not low-risk E7 reduced the steady-state levels of RB1 protein (Figure 2). Earlier studies demonstrated that low-risk E7 can bind to RB1, disrupt the interaction of E2F with RB1, and induce the expression of E2F reporter constructs to varying degrees, but they did not test the effect of low-risk E7 on endogenous E2F target genes (34–37). Here we found that E2F target gene expression was inversely related to the level of RB1 protein in HPV E7-expressing cells. High-risk HPV E7 reduced RB1 levels and caused robust activation of E2F-dependent gene expression whereas low-risk E7 had little effect on RB1 levels or E2F-dependent gene expression (Figure 2, 3). In contrast, both high-and low-risk HPV E7 reduced the steady-state levels of PTPN14, albeit to varying degrees (Figure 2). High- and low-risk E7 expression reduced the expression of epithelial differentiation markers, in both cases dependent on the C-terminal arginine that binds PTPN14 (Figure 3). With respect to their effects on RB1 and PTPN14, we conclude that the predominant difference between high- and low-risk E7 is that high-risk E7 can target RB1 for degradation. Although there are differences in the affinity of high- and low-risk E7 for PTPN14, all the E7 tested here could reduce PTPN14 levels. Based on these findings, we predicted that depletion of RB1 would cooperate with low-risk HPV6 E7 to extend the lifespan of primary keratinocytes. Indeed, RB1 depletion plus HPV6 E7 (but not HPV6 E7 R77S) extended keratinocyte lifespan (Figure 5), further supporting that inactivation of both tumor suppressors is required for keratinocyte lifespan extension by E7.

PTPN14 depletion could similarly cooperate with HPV6 E7 to extend keratinocyte lifespan (Figure 6). To gain insight into possible mechanisms underlying the long-term survival of HPV6 E7 + sgPTPN14 cells in the lifespan extension assay, we analyzed global gene expression patterns in cells expressing HPV6 E7 and/or depleted of PTPN14 (Figure 6F). Immediately after transduction, PTPN14 knockdown was associated with a modest increase in the expression of several hallmark E2F target genes. After several months in culture, the HPV6 E7 + sgPTPN14 cell pools and the HPV16 E7 cell pools exhibited increased expression of E2F target genes and decreased expression of differentiation related genes. Although PTPN14 knockdown alone was insufficient for lifespan extension, in combination with HPV6 E7 it allowed for the outgrowth of a cell population that, with respect to the expression of E2F targets and differentiation genes, resembled HPV16 E7 cells. Such a finding could be explained by reports in the literature that certain E2F target genes, including CCNE2, are subject to cooperative regulation by E2F and YAP1 (69–74). We speculate, and the data in Figure 6F support, that experimental depletion of PTPN14 can cause increased expression of E2F target genes, perhaps in a minority of cells or at a low level, and that long-term culture enables selection for and outgrowth of that population. However, PTPN14 depletion could only enable long-term cell survival in combination with low-risk E7. Our future experiments will aim to determine whether synergistic effects of E2F and YAP1 account for the increased expression of certain growth-promoting genes that could enable lifespan extension.

We propose that the findings presented here have implications for HPV biology and carcinogenesis. The E7/RB1 and E7/PTPN14 interactions are likely both important for persistent HPV infection in stratified epithelia. Here, we cultured cells in conditions that support robust proliferation, much like the cells in the proliferative basal layer of a stratified epithelium and found that low-risk HPV E7 had a minimal effect on E2F target gene expression. A possible explanation is that in proliferative cells, the baseline level of E2F target gene expression is relatively high and is not increased by low-risk E7. It is likely that in differentiated cells low-risk HPV E7 can induce E2F target expression and indeed experiments in tissue models show that low-risk HPV E7 induces expression of the E2F target *PCNA* in suprabasal, differentiated keratinocytes (10, 75). In contrast, both high- and low-risk HPV E7 proteins reduced PTPN14 levels and differentiation gene expression, suggesting that inactivation of PTPN14 is a conserved E7 function likely important for the HPV life cycle. In other work, we determined that when high-risk HPV E7 proteins degrade PTPN14, they activate YAP1, thereby inhibiting differentiation gene expression and promoting the retention of keratinocytes in the basal epithelial compartment (50). More experiments will be required to determine whether low-risk HPV E7 proteins similarly promote basal epithelial identity.

Our results could help to explain several findings in the human cancer biology literature. Germline PTPN14 mutations predispose individuals to develop basal cell carcinoma and cervical cancer (52) and somatic mutations in PTPN14 are enriched in basal cell carcinomas (53). Our data may help to explain how genetic inactivation of PTPN14 could increase carcinogenic potential both in the presence and absence of HPV infection. In addition, HPV42, previously classified as low-risk, can be carcinogenic at a specific anatomical site (76), but the level of PTPN14 expression in cells in which those cancers arise is not known. It is possible that low-risk HPV E7 has some carcinogenic potential in cells that do not express PTPN14. In the long term, it will be important to determine how PTPN14 inactivation and RB1 inactivation synergize to regulate gene expression in experimental models and to contribute to cancer development *in vivo*.

Overall, our results support the model that much of the biological activity of high- and low-risk HPV E7 proteins depends on inactivation of the RB1 and PTPN14 tumor suppressors. The degree to which E7 proteins inactivate RB1 accounts for much of the difference between high- and low-risk E7 proteins, yet PTPN14 inactivation is essential for the biological activity of both high- and low-risk E7. In the long term, our goal is to determine how the balanced regulation of RB1 and PTPN14 controls gene expression. We note that many binding partners of E7 besides RB1 and PTPN14 have been reported (13) and an additional long-term goal is to determine whether other host cell targets are required for the carcinogenic activity of certain HPV E7 proteins, for their ability to promote HPV persistent infection, or both.

## Materials and methods

### Plasmids and cloning

#### HPV E7 expression in human keratinocytes

MSCV-puro C-terminal FlagHA retroviral expression vectors encoding HPV16 E7 wild-type (WT), HPV16 E7 ΔDLYC, HPV16 R77S, HPV18 E7 WT, HPV18 E7 R84S, HPV6 E7 or GFP vector controls were previously described (21, 40) (Supplemental Table 2). Additional vectors encoding various HPV E7 were cloned by Gateway recombination of pDONR E7 plasmids with MSCV-P C Flag HA GAW, MSCV-neo-C-HA GAW or MSCV-hygro C-HA GAW destination vectors. The R84A L91A mutation was introduced by site-directed mutagenesis into pDONR 18E7-Kozak and recombined into destination vectors with neomycin selection as previously described.

#### Biolayer interferometry

All constructs were generated by standard PCR and isothermal assembly cloning methods unless otherwise noted. gBlocks encoding HPV18, HPV16, HPV11, HPV6 E7, and the C-terminal phosphatase domain of PTPN14 (residues 886–1147) were purchased from Integrated DNA Technologies. E7 constructs were cloned into a modified pET15b vector containing an N-terminal MBP-tag, His-tag, and tobacco etch virus (TEV) cleavage site (MHT-tag). PTPN14 was cloned into the N-terminal MHT-tagged pET15b vector with an additional C-terminal Twin-Strep (TS) tag. Constructs were verified by Sanger sequencing (Genewiz/Azenta Life Sciences).

#### CRISPRi

The KRAB-dCas9-MeCP2 cassette was excised from lentiCRISPRi vector (Addgene #170068) by double digestion with BamHI and AgeI and gel purified. The cassette was ligated into the corresponding BamHI/AgeI sites of plentiCRISPRv2 backbones carrying neomycin (Addgene #98292) or puromycin (Addgene #52961) selection markers to create lentiCRISPRi_Neo and lentiCRISPRi_Puro. The constructs were propagated in Stbl3 E. coli, purified by midi-prep, and verified by sequencing. Sequences for three distinct non-targeting sgRNAs were previously reported (77) and four independent sgRNAs targeting RB1 or PTPN14 promoters were chosen using CRISPick (78). The sgRNA sequences were phosphorylated and annealed, and the resulting double-stranded inserts were ligated into Esp3I-digested lentiCRISPRi_Neo or lentiCRISPRi_Puro vectors.

### Cell culture, virus production, and lifespan extension assay

Deidentified primary human foreskin keratinocytes (HFKs) were obtained from the Skin Biology and Diseases Resource-based Center (SBDRC) at the University of Pennsylvania. Cells were cultured in Keratinocyte serum free media (K-SFM, ThermoFisher) modified as previously described (12) and all experiments were performed starting with HFK at passage 4 after initial isolation. Retroviral and lentiviral particles were generated by transient transfection of HEK293T cells as previously described (12). HPV E7 expressing cell populations were established by transduction with MSCV retroviral vectors, followed by appropriate selection. RB1 and PTPN14 knockdown or nontargeting control primary HFKs were established by transduction with lentiCRISPRi vectors, followed by appropriate selection. HPV E7 expression and CRISPRi in combination was achieved by simultaneous transduction of MSCV retroviral vectors plus a lentiviral vector and combined selection with neomycin plus puromycin. Following transduction, cells were trypsinized, counted, and seeded in 10 cm dishes for selection with appropriate antibiotic for 3-5 days to establish stable populations. Once selected, cells were maintained in K-SFM and passaged every 4-5 days for up to four months. At each passage cells were counted and cell count data was used to calculate cumulative population doublings (PD). Population doublings for a single passage were calculated as PD = 3.32 (log(number of cells harvested / number of cells seeded)). Each experiment consisted of a minimum of two independent replicates conducted simultaneously unless otherwise specified.

### Western blotting

Whole cell lysates were prepared using RIPA buffer supplemented with protease and phosphatase inhibitors. The samples were sonicated to obtain homogeneous lysates. Protein concentrations were quantified using the Bradford assay. Equal amounts of protein were resolved using Mini-Protean or Criterion Tris/Glycine SDS-PAGE gels (Bio-Rad) and transferred to polyvinylidene difluoride (PVDF) membrane. Membranes were blocked with 5% non-fat dry milk in TBST (Tris buffered saline of pH 7.4 with 0.05% Tween 20) and probed with primary and HRP-conjugated secondary antibodies (Supplemental Table 3). Signals were detected by chemiluminescence.

### RNA extraction and qRT-PCR

Total cellular RNA was isolated using the NucleoSpin RNA extraction kit (Macherey-Nagel/Takara). cDNA was generated using the high-capacity cDNA reverse transcription kit (Applied Biosystems). The synthesized cDNAs were used as a template for qRT-PCR using Fast SYBR green master mix (Applied Biosystems) and a QuantStudio 3 system (ThermoFisher Scientific). KiCqStart SYBR green primers for qRT-PCR (Millipore Sigma) were used to quantify the following transcripts: PCNA, CCNE2, MCM2, RB1, KRT1, KRT10, PTPN14, and GAPDH.

### RNA-seq

Primary HFKs were transduced with retroviruses encoding GFP, untagged HPV16 E7, untagged HPV16 E7 ΔDLYC, untagged HPV6 E7, or untagged HPV6 E7 ΔGLHC. Each transduction was performed in triplicate, and independent replicates were maintained at each subsequent step. For the experiment shown in Figure 3, Supplemental Figure 4, and Supplemental Figure 5, total RNA was isolated using the RNeasy mini kit (Qiagen). PolyA selection, reverse transcription, library construction, sequencing, and initial analysis were performed by Novogene. Differentially expressed genes were selected based on a 1.5-fold change and adjusted *p-* value ≤0.05. KEGG pathways for which genes are enriched among the differentially expressed genes were identified using DAVID (79, 80). RNA-seq data have been deposited in NCBI GEO with accession number GSE319393. For the experiment shown in Figure 6F, total RNA was isolated using the NucleoSpin RNA extraction kit (Macherey-Nagel/Takara). PolyA capture, reverse transcription, second strand synthesis, tagmentation, sequencing, and initial analysis were performed by Plasmidsaurus. Differentially expressed genes were identified by comparing HPV16 E7 + sgNT (day 8) to GFP + sgNT (day 8) samples. The resulting list was compared to the Hallmark E2F Targets gene set to identify E2F-responsive genes regulated by HPV16 E7. These genes were ranked according to log_2_ fold change, and the 13 most differentially expressed genes, plus PCNA and MCM2, were selected for visualization along with keratinocyte differentiation genes previously shown to be downregulated upon PTPN14 knockout (21). The normalized expression values for the selected genes are displayed in a heatmap. RNA-seq data have been deposited in NCBI GEO with accession number GSE342734.

### Protein expression and purification

For E. *coli* expression, plasmids were transformed into E. *coli* BL21(DE3) (Invitrogen) cells, grown in Luria-Bertani (LB) medium at 37°C to an optical density of 0. 6–0.8, and induced with 0.5 mM IPTG overnight at 18°C. Cells were harvested at 5000 × *g*, lysed, and clarified at 40,000 × *g* in lysis buffer containing 20 mM Tris pH 7.5, 500 mM NaCl, 25 mM Imidazole, and 5 mM beta-mercaptoethanol (BME). All proteins were initially subjected to Ni-NTA affinity purification. Clarified lysates were applied to the resin and eluted with a linear gradient of imidazole (25 mM - 500 mM imidazole in 20 mM Tris pH 8.0, 150 mM NaCl, 5 mM BME).

#### HPV E7

Following Ni-NTA affinity purification, MHT-tagged E7 constructs were purified by anion-exchange chromatography using a Q HP column (Cytiva, 17115401). E7 was eluted from the ion-exchange column with a linear gradient of NaCl (25 mM - 1000 mM NaCl in 20 mM Tris pH 8.0, 5 mM BME. E7 was concentrated using a 30 kDa molecular weight cutoff (MWCO) Amicon centrifugal filter (Millipore) then size exclusion chromatography (SEC) on Superdex 200 (Cytiva, SD200) was used as the final purification step. The final concentration of each E7 construct was determined by measuring absorbance at 280nm and E7 was stored in 20 mM HEPES, pH 7.5, 150 mM NaCl, 1 mM TCEP at −80°C for all subsequent experiments.

#### Twin-strep-tagged PTPN14 phosphatase domain

Following Ni-NTA affinity purification, MHT-PTPN14-TS was purified by a second round of affinity purification using a MBPTrap HP (Cytiva) column and elution buffer containing 20 mM Tris, pH 7.5, 300 mM NaCl, 1% Maltose, 5 mM BME. The MHT-tag was removed with recombinant tobacco etch virus (rTEV) protease. TEV-treated samples were subjected to another round of Ni-NTA affinity purification (HisTrap FF) to separate the MHT-tag from PTPN14. PTPN14 was concentrated using a 15 kDa molecular weight cutoff (MWCO) Amicon centrifugal filter (Millipore) then size exclusion chromatography (SEC) on Superdex 200 (Cytiva, SD200) was used as the final purification step. The final concentration of PTPN14 was determined by measuring absorbance at 280nm and PTPN14 was stored in 20 mM HEPES, pH 7.5, 150 mM NaCl, 1 mM TCEP at −80°C for all subsequent experiments.

### Biolayer Interferometry

BLI assays were performed using the Gator™ Label-Free Bioanalysis instrument using Strep-Tactin-XT (ST-XT) sensor probes (Gator Bio, Palo Alto, CA, USA). The ST-XT probes were coated with purified twin-strep tagged PTPN14 phosphatase domain. A binding shift was detected after incubating the PTPN14-coated probes with different concentrations of purified E7 proteins. Buffer alone was used as reference, and MBP protein alone was used as a negative control. The dissociation constant for each protein pair was determined using the Gator One 2.7 software version 2.15.5.1221.

## Acknowledgements

We thank the members of our laboratories for helpful discussions and suggestions. This work was supported by American Cancer Society grant 131661-RSG-18-048-01-MPC to EAW, National Institutes of Health R01AI148431 and R01AI182668 to EAW, R01AI187203 to EAW and KM and R35 GM143004 to JMB. The authors acknowledge the Chemical Biology Core at the H. Lee Moffitt Cancer Center & Research Institute, which is supported in part by a National Cancer Institute (NCI) support grant (P30CA076292).

---

**Supplemental Figure 1:**
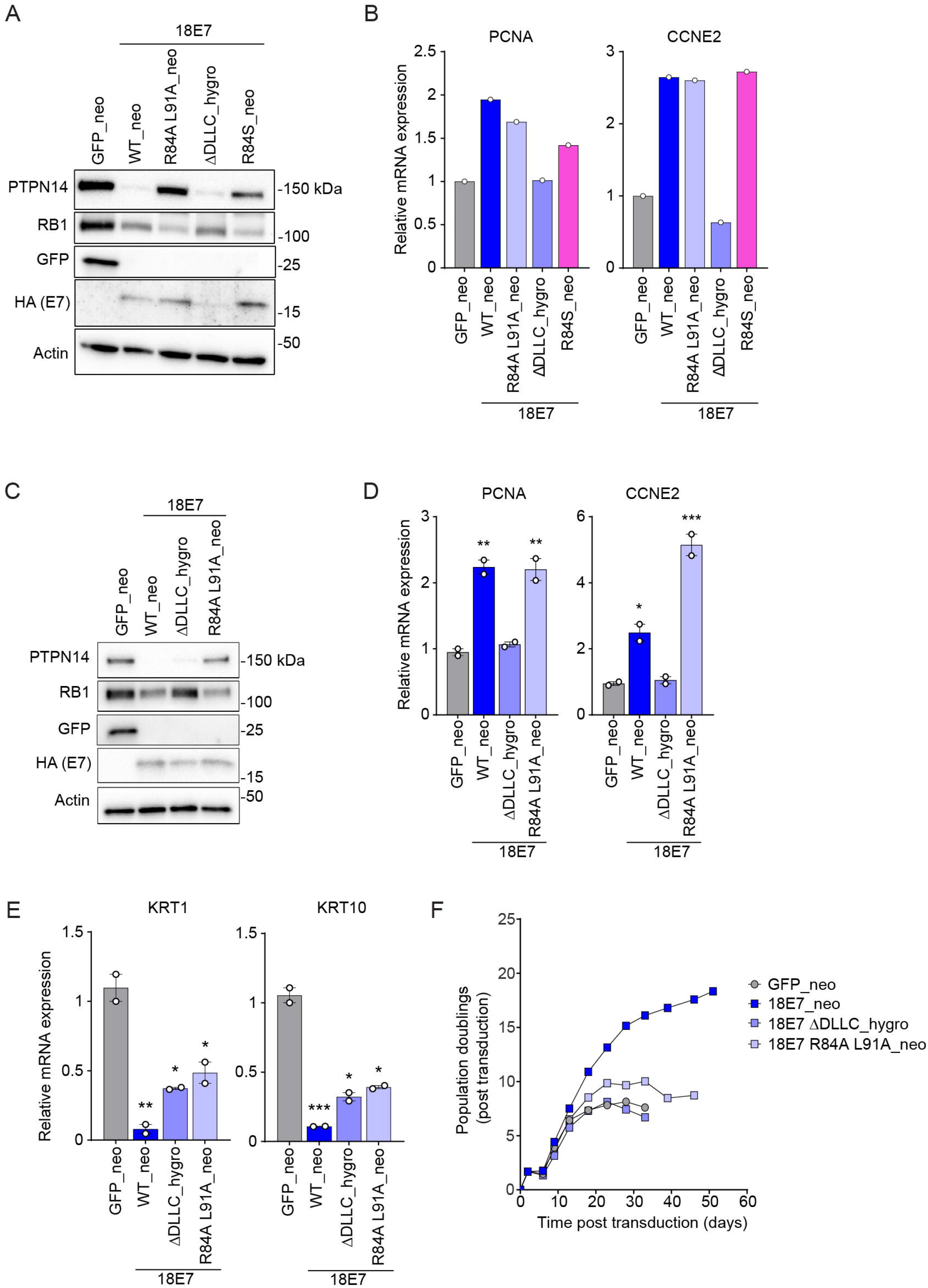
Characterization of HPV18 E7 mutants. Primary human keratinocytes were transduced with retroviral vectors encoding GFP, wild-type HPV18 E7 (WT), HPV18 E7 R84S, HPV18 E7 ΔDLLC, or HPV18 E7 R84A L91A. Retroviral vectors encoded neomycin or hygromycin resistance markers, as indicated. Each version of HPV E7 has an HA epitope tag at its C-terminus. (A) Total cell lysates were subjected to western blotting and probed with antibodies to PTPN14, RB1, GFP, HA, and actin. (B) The expression of E2F target genes (PCNA, CCNE2) was measured by qRT-PCR. (C) Total cell lysates were subjected to western blotting and probed with antibodies to PTPN14, RB1, GFP, HA, and actin. The expression of (D) E2F targets (PCNA, CCNE2) and (E) differentiation markers (KRT1, KRT10) was measured by qRT-PCR. (F) Cell populations were cultured for up to 51 days. Cells were counted at each passage and cell count data was used to calculate cumulative population doublings. In (D, E) bar graphs represent the mean ± range of two independent cell populations. Statistical significance was determined by one-way ANOVA followed by Holm-Sidak multiple comparison test, comparing each condition to the GFP control (*=*p* ≤ 0.05; **=*p* ≤ 0.01; ***=*p* ≤ 0.001). In (B, F) data points represent individual measurements from a single experiment.

**Supplemental Figure 2:**
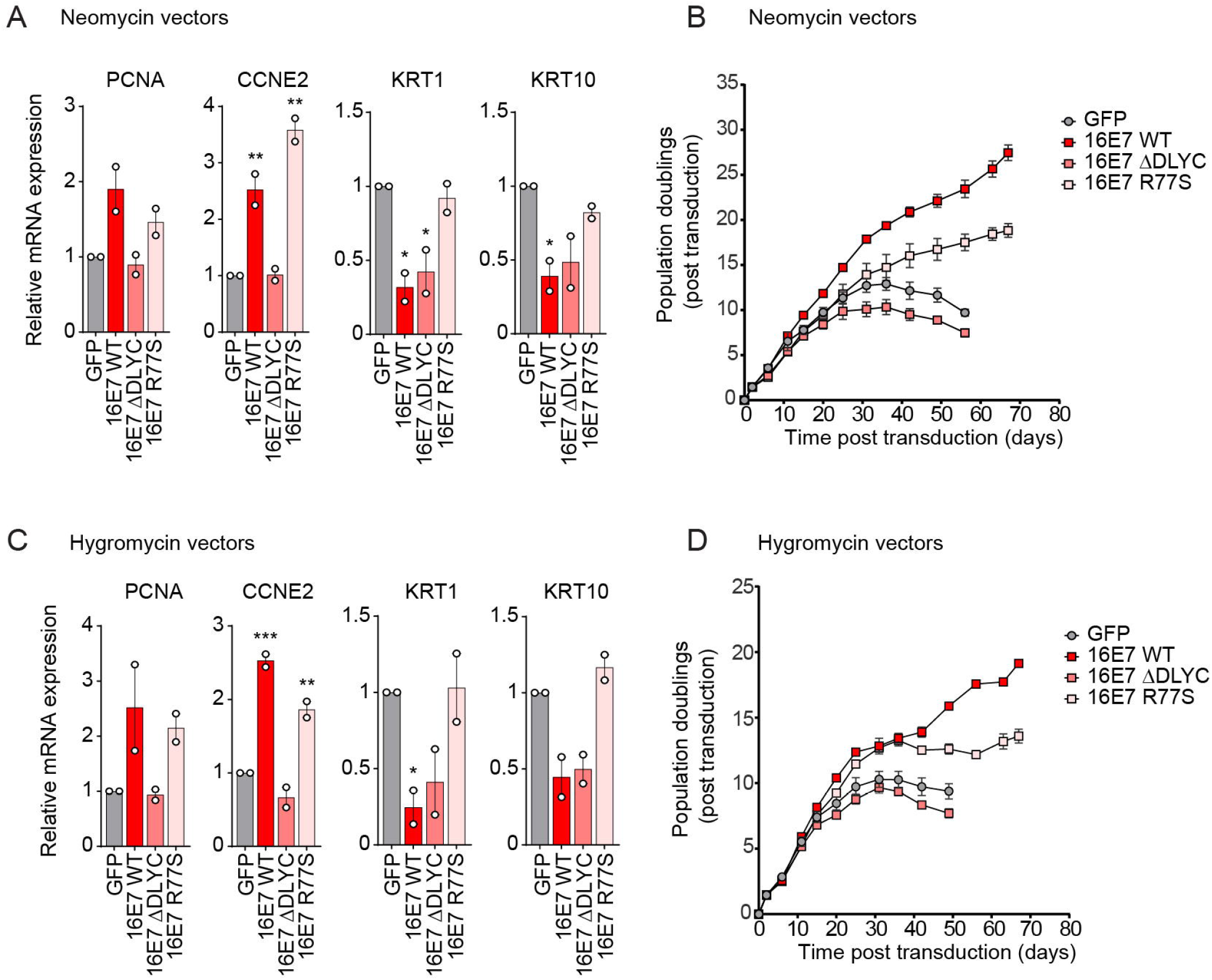
Characterization of HPV16 E7 mutants in gene expression and lifespan extension assays. (A) Primary human keratinocytes were transduced with retroviral vectors encoding GFP, wild-type HPV16 E7 (WT), HPV16 E7 ΔDLYC, or HPV16 E7 R77S. All vectors carried a neomycin resistance marker. The expression of E2F target genes (PCNA, CCNE2) and differentiation markers (KRT1, KRT10) was measured by qRT-PCR. Bar graphs represent the mean ± range of two replicate cell populations. Statistical significance was determined by one-way ANOVA followed by Holm-Sidak multiple comparison test, comparing each condition to the GFP control (*=*p* ≤ 0.05; **=*p* ≤ 0.01). (B) Cell populations from (A) were cultured for up to 67 days. Cells were counted at each passage and cell count data was used to calculate cumulative population doublings. Data points indicate the mean and error bars indicate the range for two replicate cell populations. (C) Primary human keratinocytes were transduced with retroviral vectors encoding GFP, wild-type HPV16 E7 (WT), HPV16 E7 ΔDLYC, or HPV16 E7 R77S. All vectors carried a hygromycin resistance marker. The expression of E2F target genes (PCNA, CCNE2) and differentiation markers (KRT1, KRT10) was measured by qRT-PCR. Bar graphs represent the mean ± range of two replicate cell populations. Statistical significance was determined by one-way ANOVA followed by Holm-Sidak multiple comparison test, comparing each condition to the GFP control (*=*p* ≤ 0.05; **=*p* ≤ 0.01; ***=*p* ≤ 0.001). (D) Cell populations from (C) were cultured for up to 67 days. Cells were counted at each passage and cell count data was used to calculate cumulative population doublings. Data points indicate the mean and error bars indicate the range for two replicate cell populations.

**Supplemental Figure 3:**
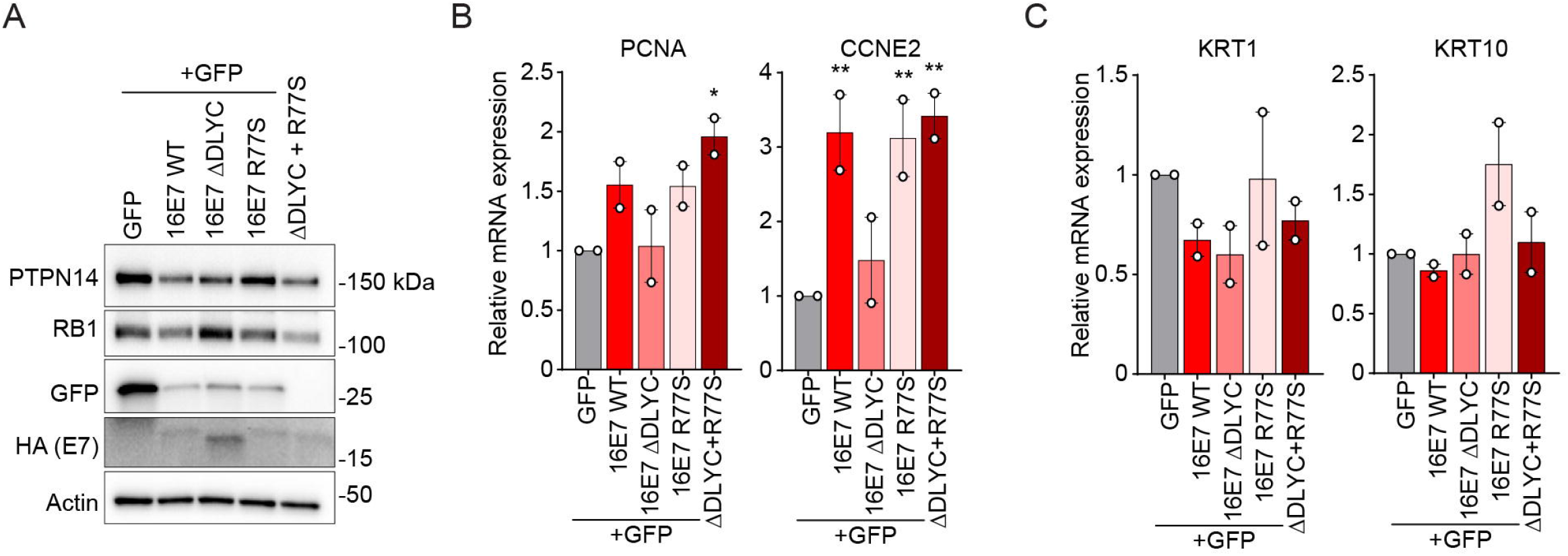
HPV16 E7 mutant proteins that cannot inactivate RB1 or PTPN14 can complement each other *in trans*. Primary human keratinocytes were transduced with pairs of retroviruses encoding GFP, wild-type HPV16 E7 (WT), HPV16 E7 ΔDLYC, or HPV16 E7 R77S, with one hygromycin-resistant and one neomycin-resistant retrovirus included in each condition. Each version of HPV16 E7 has an HA epitope tag at its C-terminus. (A) Total cell lysates were subjected to western blotting and probed with antibodies to PTPN14, RB1, GFP, HA, and actin. The expression of (B) E2F target genes (PCNA, CCNE2) and (C) differentiation markers (KRT1, KRT10) were measured by qRT-PCR. Bar graphs represent the mean ± range of two replicate cell populations. Statistical significance was determined by one-way ANOVA followed by the Holm-Sidak multiple comparison test, comparing each condition to the GFP control (*=*p* ≤ 0.05; **=*p* ≤ 0.01).

**Supplemental Figure 4.**
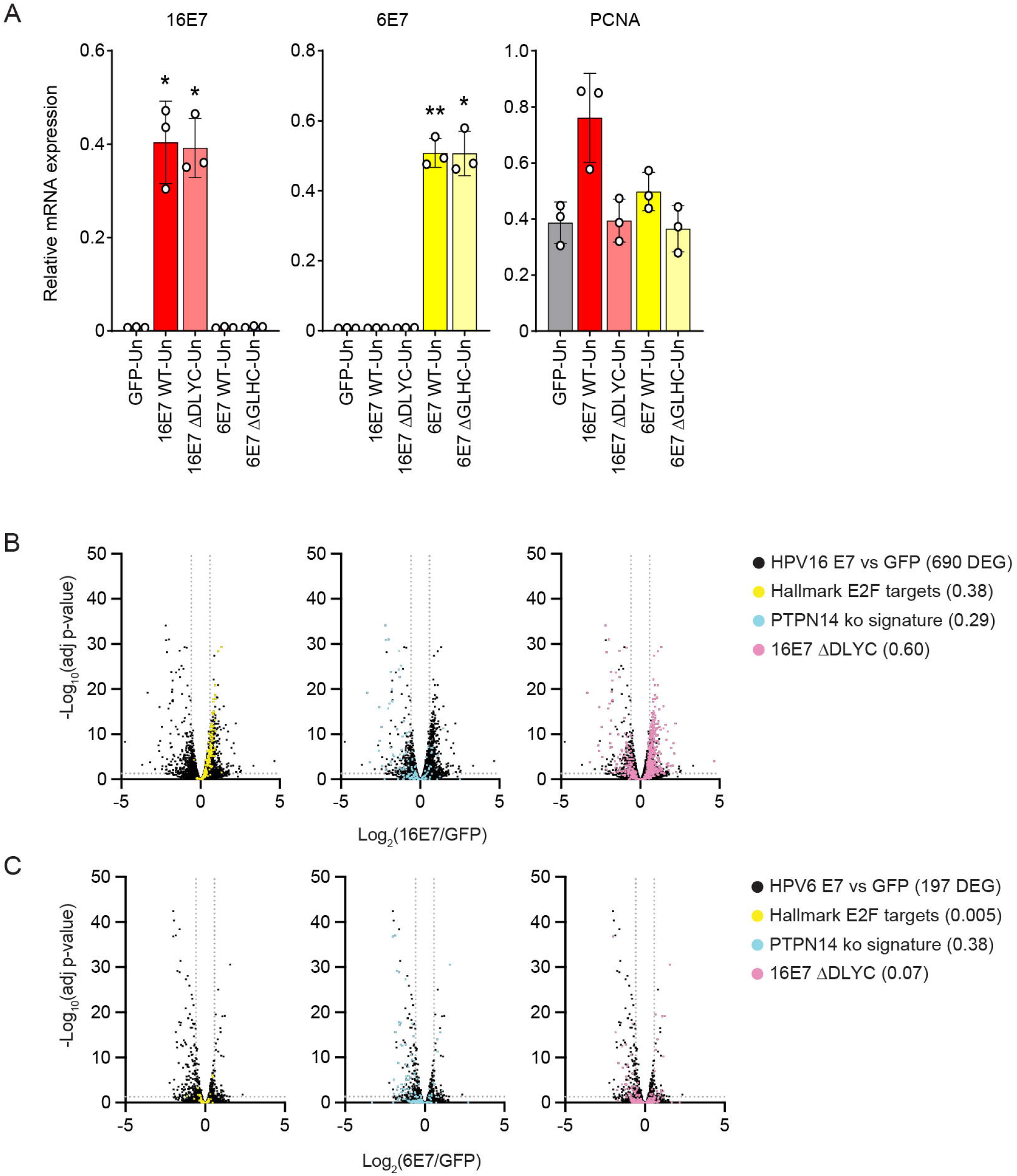
RNA-seq analysis of HFK expressing HPV16 E7 or HPV6 E7. Primary HFK were transduced with MSCV retroviral vectors encoding untagged HPV16 or HPV6 E7 proteins or GFP. (A) The expression of 16E7, 6E7, and PCNA was measured by qRT-PCR. Bar graphs represent the mean ± SD from three replicate cell pools. Statistical significance was determined by one-way ANOVA followed by Holm-Sidak multiple comparison test, comparing each condition to the GFP control (*=*p* ≤ 0.05; **=*p* ≤ 0.01). (B) PolyA selected RNA from HFKs expressing HPV16 E7 or GFP was analyzed by RNA-seq. Each volcano plot shows the gene expression changes in HFK HPV16 E7 vs HFK GFP. 690 genes are differentially expressed with ≥ 1.5-fold change and adjusted *p-*value ≤ 0.05. DEG = differentially expressed genes. (C) PolyA selected RNA from HFKs expressing HPV6 E7 or GFP was analyzed by RNA-seq. Volcano plots show gene expression changes in HFK HPV6 E7 vs HFK GFP. 197 genes are differentially expressed with ≥ 1.5-fold change and adjusted *p-*value ≤ 0.05. For (B) and (C), vertical dashed lines denote 1.5-fold change and horizontal dashed line denotes adjusted *p-*value = 0.05. In the left panel, the 200 genes that comprise the Hallmark E2F targets list are highlighted in yellow. In the center panel, the 140 genes that are differentially expressed upon PTPN14 knockout in HFKs (≥ 1.2 fold change and adjusted *p-*value ≤ 0.05) (55) are highlighted in blue. In the right panel, the 713 genes that are differentially expressed in HPV16 E7 cells relative to HPV16 E7 ΔDLYC cells in this experiment are highlighted in pink. The ratios in the figure legends indicate the fraction of genes in a list that were differentially expressed in HFK-E7 vs HFK-GFP cells in this experiment.

**Supplemental Figure 5.**
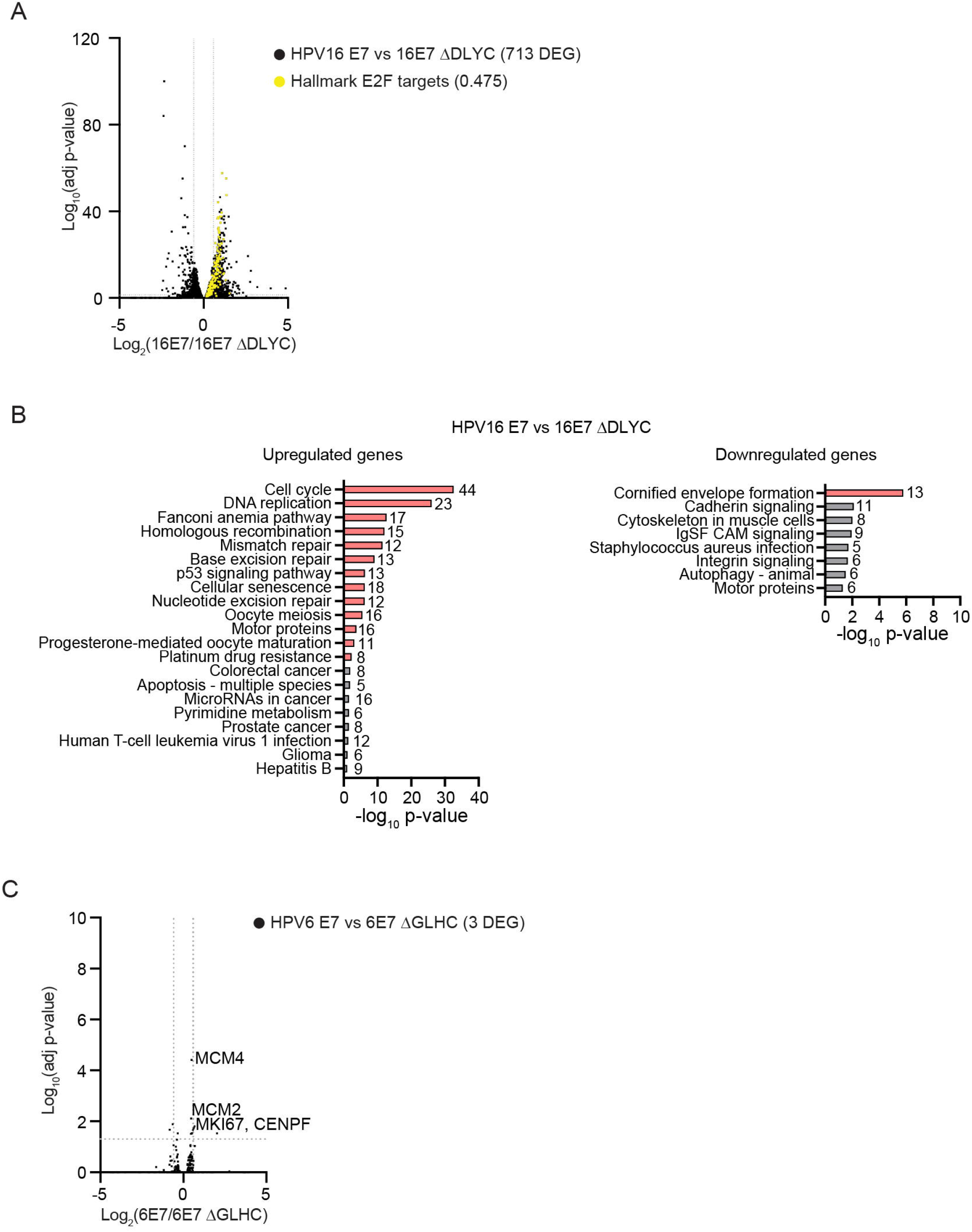
RNA-seq analysis of HFK expressing HPV16 E7 or HPV6 E7 compared to the corresponding LxCxE mutant. (A) Primary HFKs were transduced with retroviral vectors encoding untagged HPV16 E7 or HPV16 E7 ΔDLYC proteins. PolyA selected RNA from HFK expressing HPV16 E7 or HPV16 E7 ΔDLYC was analyzed by RNA-seq. The volcano plot shows the gene expression changes in HFK HPV16 E7 vs HPV16 E7 ΔDLYC. Over 700 genes are differentially expressed with ≥ 1.5-fold change and adjusted *p-*value ≤ 0.05. 47.5% of the hallmark E2F target genes are differentially expressed in HFK HPV16 E7 vs HPV16 E7 ΔDLYC. (B) Following RNA-seq, KEGG pathway enrichment analysis was conducted on the genes that were upregulated (≥1.5-fold higher expression and adjusted *p*-value ≤ 0.05) or downregulated (≥1.5-fold lower expression and adjusted *p*-value ≤ 0.05) in HPV16 E7 cells relative to 16E7 ΔDLYC cells. Pathways with *p*-values ≤ 0.05 are included in the chart. Bars for pathways with FDR ≤ 0.05 are colored whereas bars for pathways with FDR > 0.05 are in grey. Values to the right of each bar indicate the number of genes in the list associated with a pathway. (C) Primary HFKs were transduced with retroviral vectors encoding untagged HPV6 or HPV6 E7 ΔGLHC proteins. PolyA selected RNA from HFK expressing HPV6 or HPV6 E7 ΔGLHC was analyzed by RNA-seq. The volcano plot shows the gene expression changes in HFK HPV6 E7 vs HPV6 E7 ΔGLHC. Three genes are differentially expressed with ≥ 1.5-fold change and adjusted *p-*value ≤ 0.05. Selected E2F target genes are labeled.

**Supplemental Figure 6.**
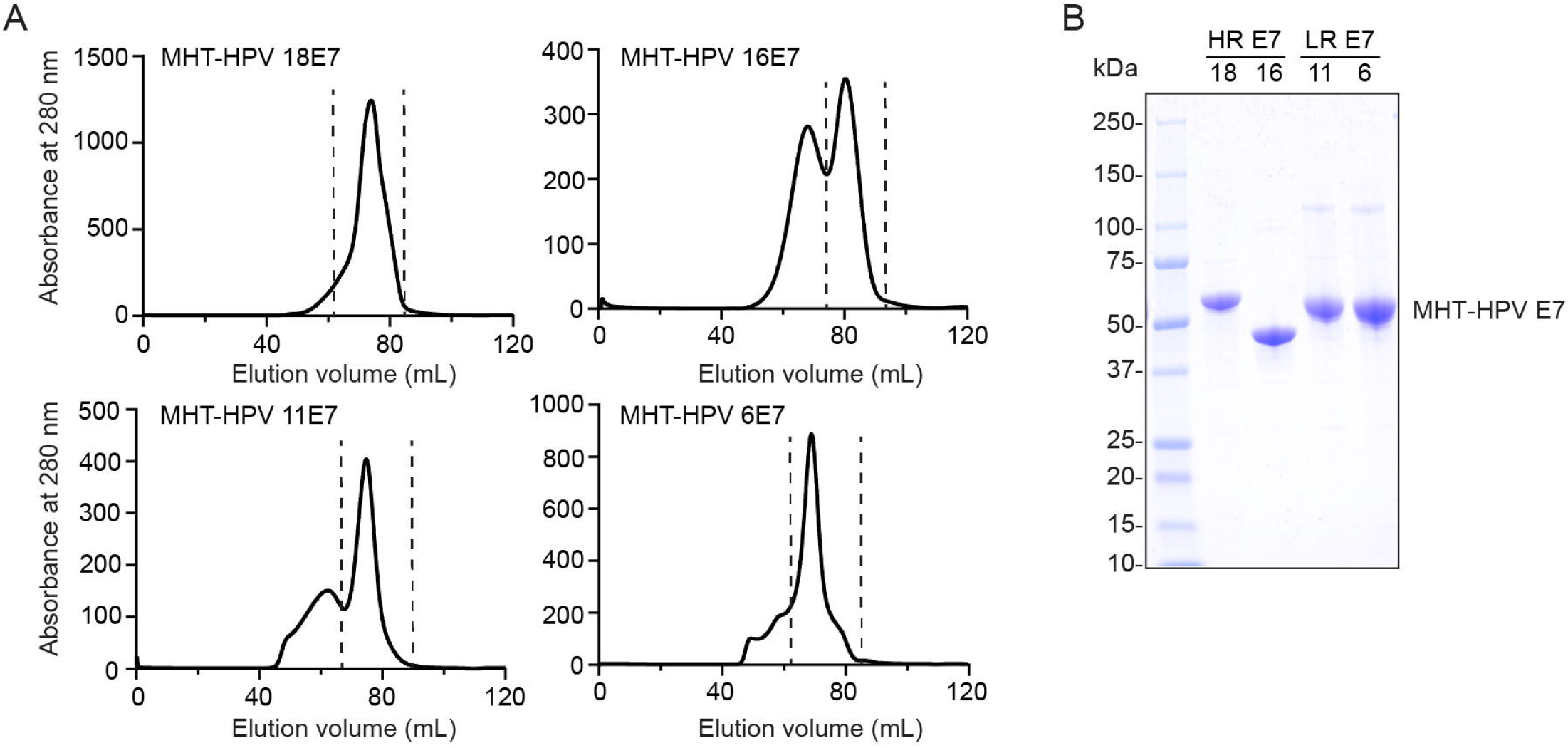
Purification of HPV E7 proteins. Recombinant MHT-E7 proteins were purified from bacteria. (A) Size exclusion chromatogram (SEC) profiles of the different E7 proteins used for affinity measurements. Fractions between the dotted lines were pooled and used for downstream BLI assays. (B) SDS-PAGE of pooled fractions for each MHT-E7.

**Supplemental Figure 7:**
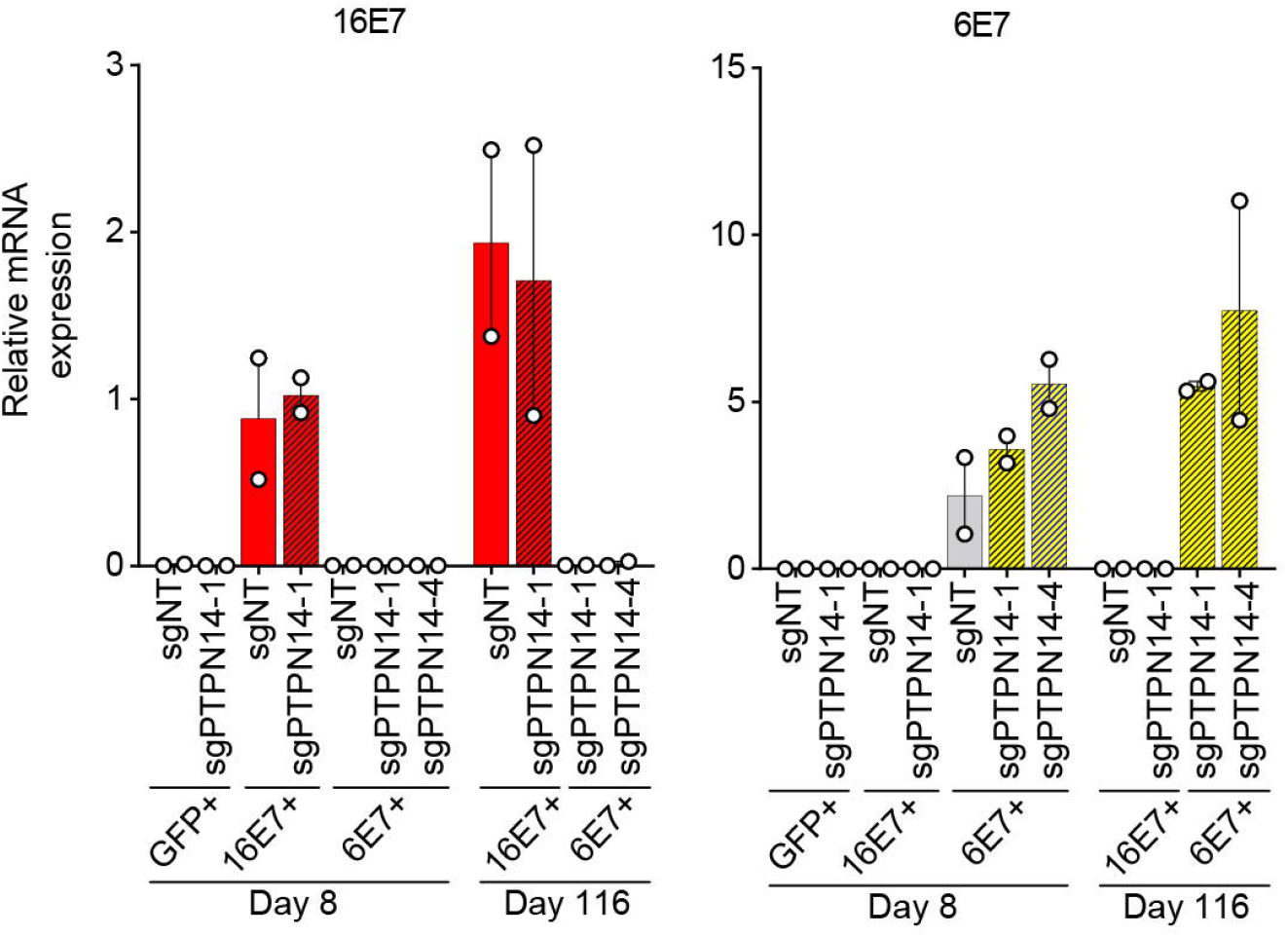
E7 expression in long-term complementation experiment with CRISPRi mediated PTPN14 knockdown. Primary human keratinocytes were transduced with a retrovirus encoding GFP, HPV16 E7, or HPV6 E7, plus a CRISPRi lentivirus and selected with puromycin and neomycin. HPV16 E7 and HPV6 E7 RNA levels were measured by qRT-PCR in the day 8 and day 116 cell populations in figure 6E. Bar graphs represent the mean ± range of two replicate cell populations.

**Supplemental Table 1. KEGG pathway analysis**

**Supplemental Table 2. Plasmids used in the study**

**Supplemental Table 3. Antibodies used in the study**

## Notes

### Competing Interest Statement

The authors have declared no competing interest.

### Summary of Updates

Updated figures 5 and 6, Supplemental files updated

